# A chromosome-scale Plasmodium cynomolgi Berok genome reveals a distinct subtelomeric architecture and a highly diverged primate malaria lineage

**DOI:** 10.64898/2026.08.21.746107

**Authors:** Adeline C.Y. Chua, Vipin Narang, Eric Jit Kai Lim, Sourav Nayak, Demi Glidden, Peter Christensen, Vaishnavi Chandramouli, Kanisha Saurin Shah, Shu Xuan Tan, Rossarin Suwanarusk, K.G. Srinivasan, Arnab Pain, Kevin Shyong Wei Tan, Peter Preiser, Bruce Russell, Georges Snounou, Laurent Rénia, Zbynek Bozdech, Bernett Teck Kwong Lee, Pablo Bifani

## Abstract

*Plasmodium cynomolgi* is the closest relative of *P. vivax* and the primary experimental model for relapsing malaria, hypnozoite biology, and blood-stage drug susceptibility. Yet existing reference genomes remain fragmented, leaving structurally complex, AT-rich regions largely unresolved. We generated a chromosome-scale genome assembly for the K4-A7 cloned line of *P. cynomolgi* Berok by combining Hi-C chromosome conformation capture, Oxford Nanopore long reads, PacBio, and Illumina sequencing. The assembly spans 14 chromosomes plus mitochondrial and apicoplast genomes, with only seven unplaced minor contigs, the fewest for any non-*P. falciparum Plasmodium* genome, and an N50 of 3.06 Mb. Critically, this hybrid strategy resolved approximately 8 Mb of extremely AT-rich (∼20% GC) sequence onto chromosomes 4, 8, and 13, anchoring what were previously unplaced or absent contigs into a continuous chromosomal framework. These subtelomere-like expansions (SLEs) constitute ∼26.5% of the chromosomal genome and are enriched for PIR/VIR, STP1, variable surface antigen, and methyltransferase pseudogene families. Despite low gene density, SLE-encoded genes are transcriptionally active and show stage-specific expression across the erythrocytic cycle. Integrated lifecycle transcriptomics across 7,006 genes revealed a ∼54-hour erythrocytic cycle with a “just-in-time” transcriptional cascade closely resembling that of *P. vivax*. Phylogenomic analyses and pairwise amino acid comparisons across more than 2,600 single-copy orthologs show that Berok forms a deeply diverged *P. cynomolgi* lineage, suggesting a distinct subspecies. This assembly establishes a high-resolution genomic foundation for comparative malaria biology, drug discovery, and the study of subtelomeric architecture, host adaptation, and lineage boundaries in primate *Plasmodium*.

## Introduction

Human malaria is caused by a set of polyphyletic *Plasmodium* species that belong to two evolutionary clades: *Laverania* that is solely represented by *P. falciparum*^1^ responsible for the majority of recorded cases and mortality, while the other species fall into the *Plasmodium* clade. Of these, the globally distributed *P. vivax* is the second most important species. Several biological characteristics make the control of *P. vivax* challenging^2^, of which the ability to form dormant liver stages, hypnozoites, is the most important. Hypnozoites can activate to lead to relapse episodes over the months and years following the infectious mosquito bite, effectively extend the duration of the infection and the potential for further transmission. These inaccessible, and thus, little studied forms can only be eliminated by two 8-aminoquinoline drugs (primaquine and tafenoquine) whose use is restricted by potential toxicity. Phylogenetically *P. vivax* clusters with a set of parasite species that naturally infect Southeast Asian macaques^3^, with *P. cynomolgi* being the most closely related genetically and biologically and one of the three species that can form hypnozoites. The long-standing inability to date to adapt *P. vivax* to continuous *in vitro* cultivation has made *P. cynomolgi* infections of macaques an extensively used experimental model for *P. vivax*, and in particular for the search of drugs against hypnozoites^4^. The recent development of a protocol that allows the *in vitro* cultivation of the hepatic stages of *P. cynomolgi* brought the potential to investigate the hypnozoite^5^.

Accidental infections revealed that humans are susceptible to *P. cynomolgi*^6, 7^. and recently cases of naturally acquired zoonotic *P. cynomolgi* infections have been recorded in Southeast Asia^8, 9, 10, 11, 12^, though much fewer than those due to *P. knowlesi*. Although *in vitro* cultivation of *P. cynomolgi* was reported in the 1980s^13, 14^, a protocol for continuous cultures was only recently developed for the Berok strain of *P. cynomolgi*^15, 16^. This has already provided the possibility to genetic modifications and drug susceptibility investigations^17, 18^, and opens the way to conduct biological, functional, and immunological studies hitherto restricted to experimental macaque infections. In this context the availability of the complete genome will substantially enhance such studies. Moreover, this could help understand the zoonotic potential and limitations of the *Plasmodium* species in Southeast Asian macaques.

An initial genome assembly of a *P. cynomolgi* strain^19^ was later refined^20^ and provided confirmation of the close phylogenetic relationship with *P. vivax*. Although the first strain sequenced was given as *P. cynomolgi bastianellii*, denoted strain B^19^, the doubt expressed in that publication that it might have been strain M was later confirmed^20, 21^, and it will be denoted as M or so-called B in the current publication. The initial assembly based on Sanger, Roche 454 and Illumina sequencing yielded a 26,18 Mb that included 1648 unplaced contigs totalling 3,45 Mb. This was refined in the second assembly based on Pacific Biosciences SMRT technologies to yield a total genome of 30,66 Mb, including only 40 unplaced contigs that still comprised 4,7 Mb, distributed over 14 chromosomes with a predicted total of 6632 genes.

The advent of novel sequencing strategies and technologies provides a means to substantially improve the quality and completion of the assemblies. Thus, Chromosome conformation capture (Hi-C) and long-read sequencing have each proved effective for finishing *Plasmodium* genomes ^22, 23, 24, 25, 26^. Hi-C contact maps have been used to scaffold *P. falciparum* chromosomes and to resolve three-dimensional chromatin organisation in both *P. falciparum* and *P. vivax*, while integration of PacBio long reads with Hi-C data in *P. knowlesi* produced an assembly that resolved complex subtelomeric architectures and improved annotation of the *SICAvar* multigene family. In each of these cases, the combination of long reads to span repetitive regions and Hi-C contact information to order and orient contigs was essential for resolving subtelomeric regions that short reads alone could not assemble. In the context of *Plasmodium* subtelomeres, which are compositionally extreme and repeat-dense, Oxford Nanopore Technology (ONT) is particularly suited to spanning these regions: long reads >10 kb routinely bridge repeat arrays that fragment short-read assemblies, and the continuous signal data enables base-calling accuracy sufficient for variant detection in AT-biased sequence. For *P. cynomolgi*, where the subtelomeres harbour large multigene families and display growing evidence of inter-strain structural diversity, a hybrid approach combining ONT, Illumina, PacBio, and Hi-C data should deliver an accurate genome suitable for future robust comparative analyses.

Here, we apply this quadruple-platform strategy to a cloned line (K4-A7) of *P. cynomolgi* Berok adapted to continuous *in vitro* culture, to generate a chromosome-scale reference genome that resolves previously unplaced contigs, defines the AT-rich genomic compartment, and provides new insight into the transcriptional landscape and phylogeny of this distinctive *P. vivax* surrogate model. Our main goal is to characterize the structure and function of this subtelomere-like compartment including its composition, 3D organization, and role in the transcriptional program, and to place it within a core genome phylogenomic framework that quantifies the divergence of Berok from other *P. cynomolgi* strains. We set out to define the structure and content of this subtelomere-like compartment, examine its position in the three-dimensional genome, and determine whether genes in these regions are transcriptionally active during the intraerythrocytic cycle. We also compare Berok with other *P. cynomolgi* strains using a core-genome phylogenomic analysis to assess the extent of its divergence. By anchoring extreme subtelomeric sequences and analysing their activity across a 54-hour intraerythrocytic cycle, we link fine-scale genome architecture to broader questions of host adaptation and lineage diversification in primate malaria. Together, these analyses allow us to relate the newly assembled subtelomeric regions to gene regulation and to the evolutionary position of the Berok lineage within primate malaria parasites.

## Results

### A chromosome-scale *P. cynomolgi* Berok genome with minimal unplaced sequence

The Berok strain is the only *P. cynomolgi* that has been successfully adapted to continuous *in vitro* culture, providing a unique opportunity for selecting for clones and providing sufficient nucleic acid material for sequencing. Genomic DNA from mixed-stage continuous culture of *P. cynomolgi* cloned line ^15^, was sequenced across four complementary platforms comprising Hi-C, Oxford Nanopore Technology (long reads >1 kb), Illumina TruSeq paired-end (>40-fold coverage), and PacBio, and used to generate a hybrid genome assembly. Long Nanopore reads provided the initial assembly scaffold, which was then refined through iterative polishing with Nanopore and Illumina data; Hi-C–guided scaffolding was subsequently applied to organize contigs into chromosome-scale sequences that closely represent the true genome (see Materials & Methods for hybrid genome assembly). This hybrid assembly approach resulted in a highly comprehensive assembly of 14 nuclear chromosomes, along with the mitochondrial and apicoplast genomic elements. The total size of this assembly accounted for 37 Mb, leaving only 7 unplaced contigs of 67 kb genomic sequences (Table 1). This assembly currently represents one of the most complete genomic sequences among the *Plasmodium* species, likely owing to the hybrid assembly approach using data from four sequencing types. Indeed, the scaffold N50 of 3.06 Mb represents a near-doubling compared to prior *P. cynomolgi* assemblies (B strain: 1.72 Mb; M strain: 1.72 Mb). The three *P. cynomolgi* genome assemblies were highly complete in the BUSCO analysis with the Berok strain being the most complete (n = 4,203), 97.2% complete, with 0.7% fragmented and 2.1% missing in Berok, 95.5% complete, 2.3% fragmented, and 2.2% missing in M strain, and 90.5% complete, 7.2% fragmented, and 2.3% missing in the so-called B strain, indicating well-represented gene space across assemblies.

**Table 1.** Assembly quality metrics for non-*Plasmodium* genomes. The assembly statistics for the *P. cynomolgi* Berok K4-A7 cloned line compared to that of other *P. cynomolgi* strains as well as *P. vivax* and *P. knowlesi*. *P. cynomolgi* Berok K4-A7 has only 7 unplaced short contigs.

| Organism | Accession | No. of chromosomes | No. of unplaced contigs | Chromosome size (bp) | Unplaced contig size (bp) | Apicoplast size (bp) | Mitochondria size (bp) | Total size (bp) | GC% | N50 |
| --- | --- | --- | --- | --- | --- | --- | --- | --- | --- | --- |
| *Plasmodium cynomolgi* strain Berok |  | 14 | 7 | 36,898,196 | 67,006 | 34,480 | 5,991 | 37,005,673 | 34.50 | 3,057,228 |
| *Plasmodium cynomolgi* strain (B) M | GCF_000321355.1 | 14 | 1649 | 22,728,335 | 3,453,008 | - | - | 26,181,343 | 39.08 | 1,717,921 |
| *Plasmodium cynomolgi* strain M | GCA_900180395.1 | 14 | 40 | 25,916,758 | 4,700,316 | 34,521 | 6,017 | 30,657,612 | 37.31 | 1,715,570 |
| *Plasmodium knowlesi* strain H | GCF_000006355.2 | 14 | 148 | 23,938,832 | 420,552 | 30,638 | 5,957 | 24,395,979 | 38.60 | 2,162,603 |
| *Plasmodium vivax* strain PvP01 | GCA_900093555.2 | 14 | 226 | 24,214,674 | 4,789,968 | 29,582 | 5,989 | 29,040,213 | 39.59 | 1,761,288 |
| *Plasmodium vivax* strain Salvador I | GCF_000002415.2 | 14 | 2733 | 22,621,071 | 4,386,630 | - | 5,990 | 27,013,691 | 42.19 | 1,678,596 |

### Multi-platform sequencing confirms uniform coverage across the assembly as well as subtelomere-like expansions (SLEs)

Coverage histograms across all 14 chromosomes from five independent Illumina samples (one parental sample and four K4-A7 clonal samples derived by serial dilution ^15^; and two long-read samples from a K4-A7 clonal sample (ONT and PacBio) show high, uniform read depth across the entire assembly (Fig. 1A). This cross-sample and cross-platform concordance confirms the accuracy of the assembly approach. Nevertheless, we wish to further substantiate the overall assembly’s structural integrity and chromosomal-scale contiguity by the Hi-C contact map. This approach takes advantage of the fact that genomic loci interact more frequently in *cis* than in *trans*, and thus correctly scaffolded genomes yield high-frequency contacts along the matrix diagonal, forming distinct chromosome-specific square blocks. In agreement with this, the assembled contact map of the *P. cynomolgi* genome displays sharp, well-resolved diagonal blocks for all 14 chromosomes (Fig. 1B). This confirms accurate contig ordering and orientation with no detectable large-scale mis-assemblies. Beyond, local contiguity, the map reveals distinct A/B chromatin compartments across all chromosomes (Fig. 1C). The successful recovery of these presumably euchromatin (compartment A) and heterochromatin (compartment B) domains provides an independent biological validation. Crucially, minimal signal is observed in off-diagonal regions, indicating a high ratio of *cis*-to-*trans* interactions. This low inter-chromosomal noise indicates that the scaffolding introduced minimum chimeric artifacts or cross-chromosomal mis-joins and that the vast majority of the assembly truly reflects the organization of the *P. cynomolgi* genome. The well-defined diagonal architecture and coherent compartment structures offer robust, sequence-independent validation, confirming that chromosomal-scale ordering and orientation are correct across all 14 chromosomes. This is a form of validation that N50 and BUSCO scores alone cannot provide. An interactive Hi-C contact map is available at https://tinyurl.com/22fkpjtl. Synteny mapping between the generated genome information of the Berok strain and the reference genome of the *P. cynomolgi* M strain revealed a good agreement with the chromosomal placements of the assembled contigs across the 14 chromosomes except for chromosome 12, which appears to carry one intrachromosomal non-syntenic segment (Fig. 1D). Specifically, 22.4 Mb out of 25.9 Mb of the *P. cynomolgi* M strain chromosomal sequence aligned to the Berok strain as determined by pairwise alignments using MUMmer^27^. This shows a high degree of concordance between the two assemblies for most of the core genome. However, there are several considerable exceptions to this concordance, including an intrachromosomal non-syntenic segment on chromosome 12, and three large 5’ end extensions of chromosomes 4, 8, and 13. To the latter, we referred to as Berok-specific subtelomere-like expansions (SLEs).

**Figure 1.**
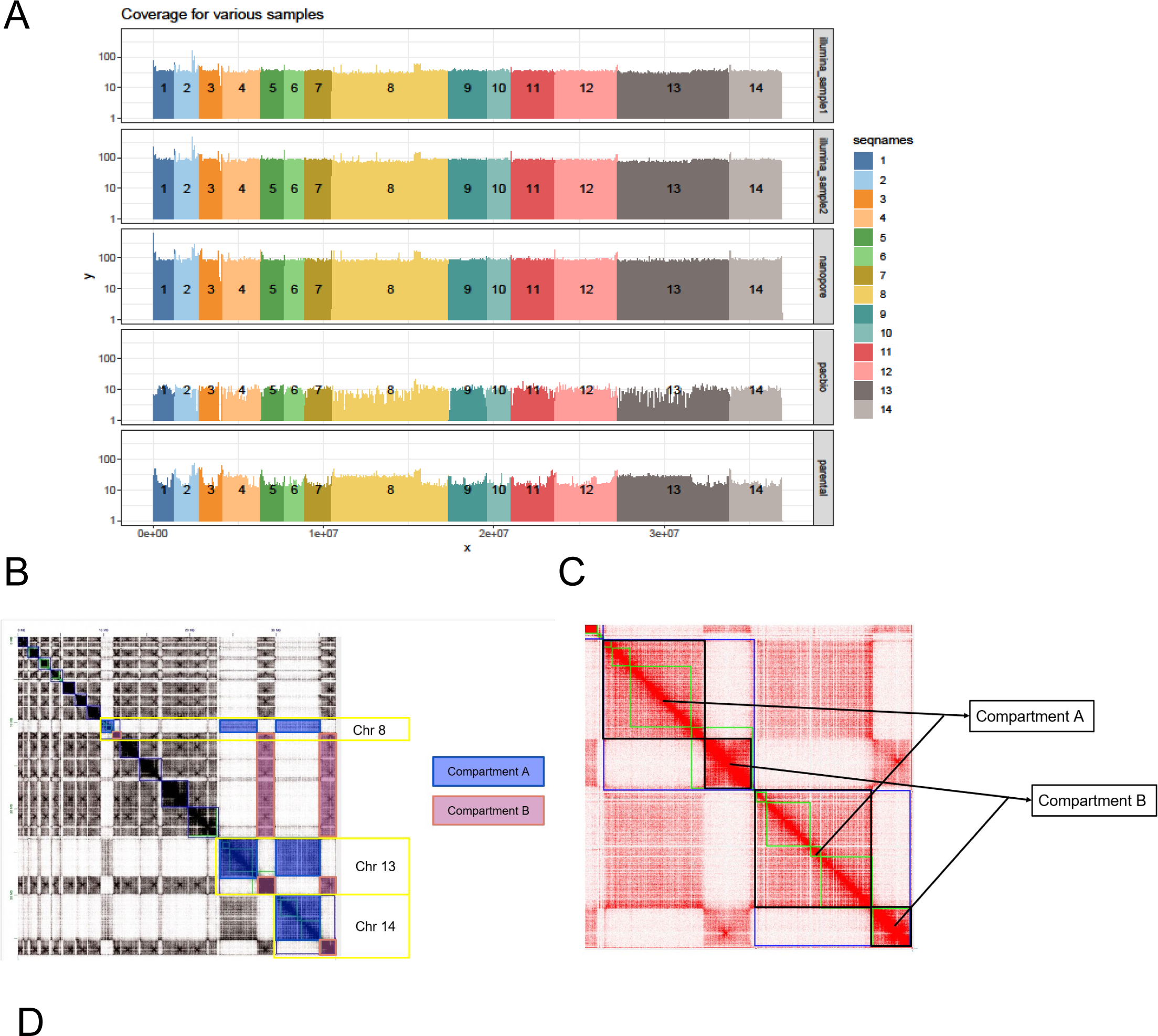
Read depth coverage across the genome for multiple independent samples derived from *P. cynomolgi* Berok. (A) Coverage histogram of multiple independent samples using both Illumina and Oxford Nanopore sequencing technology showing very high coverage over the entire *P. cynomolgi* Berok assembly. The read depth is shown in the y axis with the assembly chromosomes arranged linearly on the x axis. The histograms are colored and labelled by chromosomes. The top four samples are from Illumina sequencing while the bottom three samples are Oxford Nanopore and PacBio sequencing. (B) Cross-chromosome A/B compartmentalization across genome, most easily observed in chromosomes 8, 13, 14, but can be observed across the genome. For example, the majority of chromosomes 1-7 and 9-12 is associated with Compartment B, with small regions associated with Compartment A (see end of chromosome 11). An interactive diagram is provided URL link for Hi-C viewer (Juicebox) https://tinyurl.com/22fkpjtl. (C) Close up of Chromosomes 13 and 14 to clearly show cross-chromosome compartmentalization. (D) The chromosomes and contigs in the Berok are mapped to *P. cynomolgi* M using MUMmer alignments (colored segment). Gray segments show regions of the Berok strain which do not align. Colored segments are colored by the chromosomes on strain M. Large fragments from chromosomes 4, 8 and 13 have alignments with strain M unplaced contigs.

### Synteny mapping with the *P. cynomolgi* M reference genome reveals large subtelomere- like expansions (SLEs)

The Berok-specific SLE sequences show a sequence similarity with unplaced contigs of strain M and so-called B (black coloured regions in Fig. 1D for M strain, and Supplementary Fig. 1 for M and so-called B strain). This suggests that these expansions have been previously observed but were not placed in the previous *P. cynomolgi* genome assembly efforts^19, 20^. The sizes of the three SLEs appear to exceed the assembled portion of their aligned intrachromosomal regions, comprising 57.1%, 75.7%, and 58.2% of the core portions of chromosomes 4, 8, and 13, respectively. Overall, the SLEs (detailed in Supplementary Table 1) account for approximately 15.7 Mb (∼30% of the chromosomal genome), exhibit markedly reduced GC content (∼20% versus ∼34-42% in the remaining chromosomal regions, Supplementary Fig. 2), and show a substantially lower gene density, highlighting them as compositionally and structurally distinct from the canonical chromosomal backbone indicating that they differ markedly from the rest of the chromosomal sequence in both base composition and gene content. Contamination analysis using Kraken2 and Centrifuge further confirmed that the SLEs originates from *Plasmodium* and are not attributable to host or environmental contamination (Supplementary Fig. 3). Read coverage from four independent Illumina samples and three ONT/PacBio samples is also uniformly high throughout (Fig. 1A), including AT-rich regions in the SLEs where Illumina coverage is lower than ONT/PacBio, consistent with the known mapping bias of short reads in highly repetitive, low-complexity sequences. This further demonstrates that SLE sequences are genuine components of the Berok genome rather than assembly artifacts. The use of the Hi-C data was instrumental in identifying the positions of these expansions which showed the initial contigs prior to final assembly by Hi-C, thereby defining a previously unresolved genomic compartment in the Berok genome (Supplementary Fig. 1). Taken together, we observed significant similarities in the genomic composition of the Berok to the reference *P. cynomolgi* M and so-called B strains represented by syntenic intrachromosomal regions within all 14 chromosomes. However, evidence from multiple sequencing technologies and independent samples shows that the Berok strain genome carries at least three SLEs on chromosomes 4, 8, and 13, each with substantially different sequence characteristics in GC content and gene counts.

### Annotation concordance across *Plasmodium* species

Comparative annotation, supported by RNA-seq evidence (see below), showed that the majority of Berok strain transcript annotations have identifiable orthologs in *P. cynomolgi* M (highest proportion), followed by *P. vivax* PvP01, *P. vivax* Salvador I, *P. cynomolgi* B (M), and *P. knowlesi* H. Overall, 90% of the 5270 genes annotation in the *P. cynomolgi* Berok assembly were also detected in other *Plasmodium* species (orthology ranging 86.01% to 96.10%) (Fig. 2A). Conversely, approximately 80% of transcript annotations from these reference genomes (ranging 76.57% to 86.62%) are found in the Berok strain assembly (Fig. 2B). The genomic distribution of these identified orthologs in other *Plasmodium* species and strains (Fig. 2C) are consistent, showing that the core genome of *P. cynomolgi*, including the Berok strain, is characterized by a high and even gene density. The chromosomal locations of the identified orthologs are also highly conserved (Fig. 2D), and the gene-level synteny plot in Fig. 2E further shows that the relative positions on the chromosomes are highly conserved, with the single exception on chromosome 2, where a single synteny block appears shifted. The gene-level synteny conservation extends to *P. cynomolgi* B and *P. vivax* P01 (Supplementary Fig. 4).

**Figure 2.**
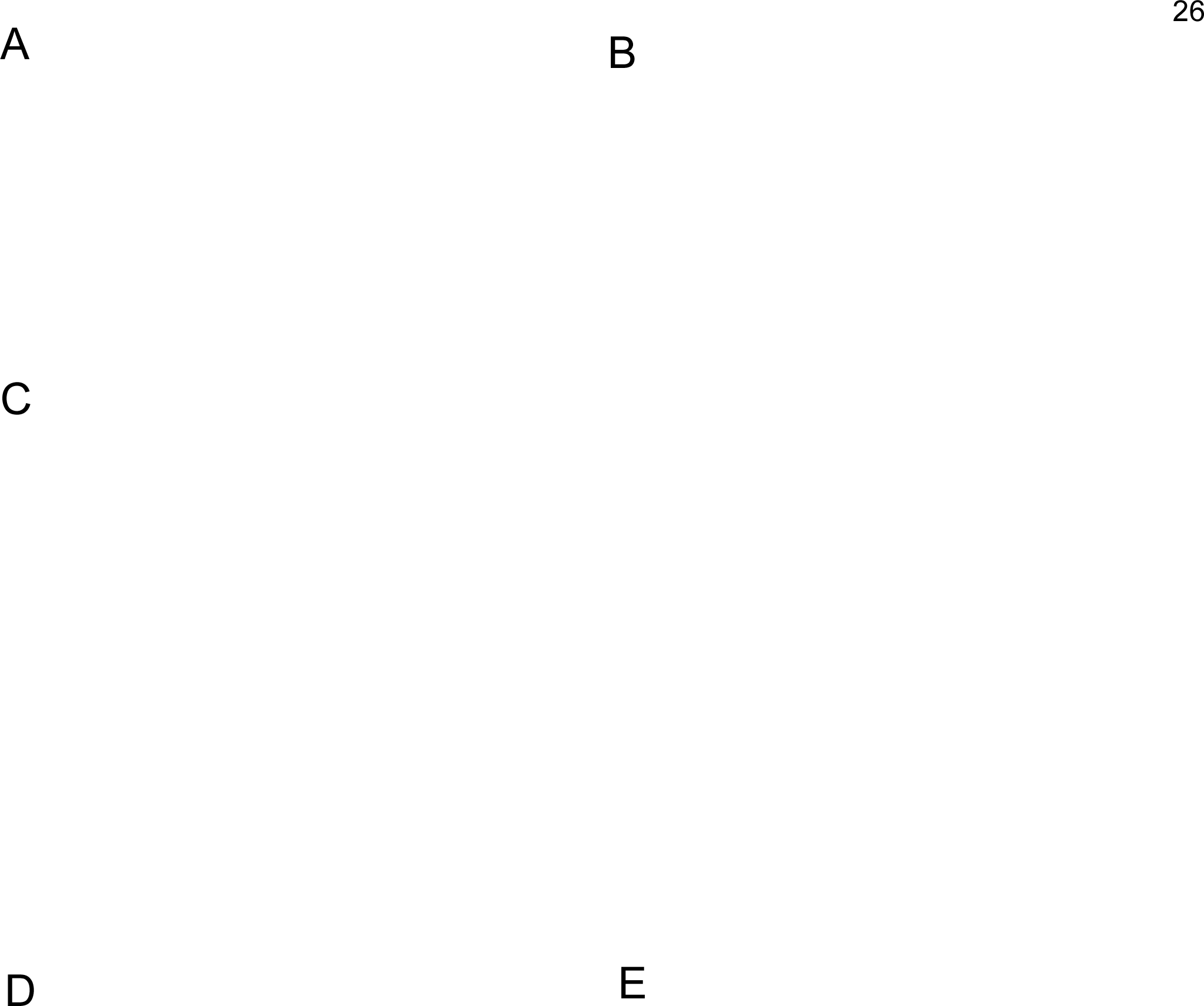
Annotation concordance across *Plasmodium* species. *P. cynomolgi* Berok annotations show good concordance with other related species including *P. cynomolgi* strain M, *P. cynomolgi* B, *P. vivax* Salvador I (Sal1), *P. vivax* PvP01 and *P. knowlesi* H. Majority of the transcript annotations in the Berok are found in *P. cynomolgi* M followed by *P. vivax* PvP01 as showed in (A). The reverse is also true as shown in (B) with about 80% of the transcript annotations in the other genomes found in the Berok strain annotations. (C) shows that the Berok strain annotations are matched throughout the genome (even for chromosomes 8 and 13). The matched genes are consistent with regards to the chromosome that they reside on as shown in the heat map in (D) where >90% of the genes are conserved in terms of chromosome location against *P. cynomolgi* M. The conservation is further highlighted in (E) showing the synteny of the genes against *P. cynomolgi* M.

Codon usage analysis (Supplementary Fig. 5A) confirms that the Berok strain genes follow overall *Plasmodium* codon usage patterns, suggesting they are subjected to similar translational selection constraints as core genome genes. However, SLE genes do differ in their codon usage (Supplementary Fig. 5B). Analysis of k-mer composition (Supplementary Fig. 6) reveals enrichment of specific 9-mers in the extended regions relative to the common regions, consistent with the highly repetitive, AT-biased sequence content of the SLEs.

### Transcriptional cascade of the *P. cynomolgi* Berok intraerythrocytic developmental cycle (IDC) shares similarities and differences with *P. vivax*

In the next step of the analysis, we sought to reconstruct the transcriptional cascade of *P. cynomolgi* Berok parasites to further validate our gene predictions and to explore the gene expression pattern of this parasite line throughout its asexual blood-stage development. For that we cultured highly synchronized *P. cynomolgi* Berok K4-A7 parasites in macaque erythrocytes as previously described^15^. From this parasite culture, cell samples were collected every 6 hours over 54 hours for RNA-seq analysis. The resulting transcriptome comprised 10 experimental time points, each presumably representing the transcriptional profile of the corresponding temporal state of *P. cynomolgi* IDC (Supplementary Table 2). Principal component analysis revealed a gradual gene expression in each time point collectively constituting a complete IDC transcriptional cascade with a duration of 54 hours (Fig. 3A). For the overview of the *P. cynomolgi* Berok IDC transcriptional cascade, we plotted smooth expression profiles for each of the detected 4,249 genes according to their periodic temporal expression characteristics using Fourier transformation as previously described^28^. Indeed, ranked by the time of their maximum expression, the *P. cynomolgi* Berok IDC transcriptional cascade forms a clear wave-like pattern that moves from early to late stages of the parasites, mirroring the “just in time” transcriptional program described in *P. falciparum* and *P. vivax* (Fig. 3B). Comparing *P. cynomolgi* to *ex vivo P. vivax*^28^ data showed that for most orthologous genes, their periodic pattern and peak expression were analogous between the two *Plasmodium* species. However, the timing of expression for a set of 944 genes differed (Fig. 3C and D). This phase shifted genes are enriched for translation, proteolysis, and central metabolic pathways, and include many ribosomal, cytosolic, and nuclear components (Fig. 3E). Taken together, these results are consistent with previous studies showing that while most genes exhibit constant expression patterns, ∼15% of the IDC transcriptional cascades vary between *Plasmodium* parasite species, likely representing their unique evolutionary adaptations^29^.

**Figure 3.**
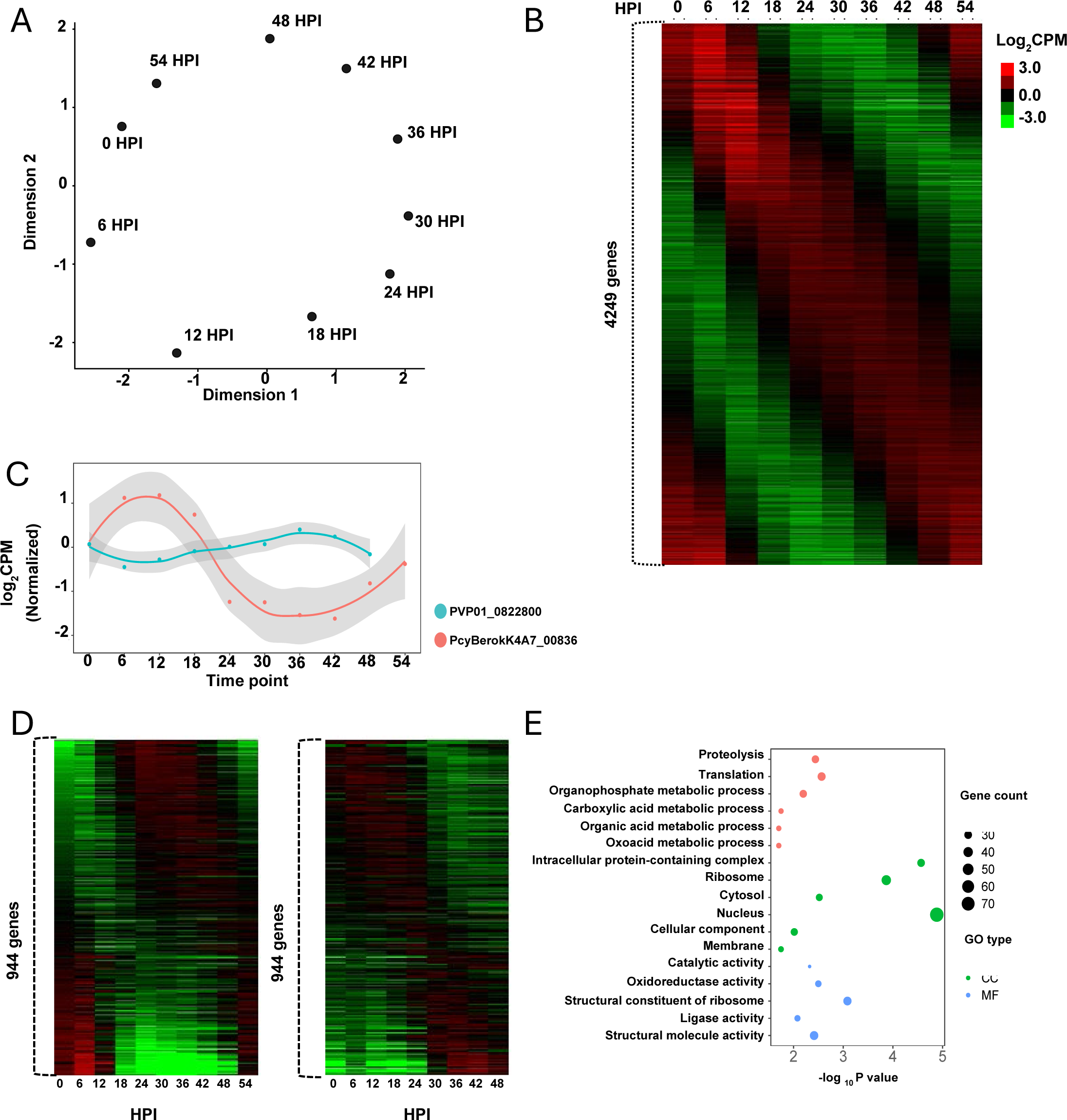
Transcriptome of *Plasmodium cynomolgi* Berok strain. (A) Multi-dimensional plot showing first two dimensions generated using TMM-normalized expression of genes during IDC time-points. (B) Expression of genes for IDC time-points showing ‘just-in-time’ transcriptional pattern for *Plasmodium cynomolgi* Berok strain. (C) Orthologous gene expression pattern of *Plasmodium cynomolgi* Berok strain and *ex vivo* isolates of *Plasmodium vivax* were compared. An example of orthologous pair of genes between these two species is shown with peak expression at different timepoints depicting differences in transcriptional cascade. (D) A total of 944 genes found two have phase difference > 2 and shown as a heatmap *Plasmodium cynomolgi* Berok strain (left) and *Plasmodium vivax* (right). (E) Enrichment based on 944 *Plasmodium vivax* geneids highlighting GO terms involving several Biological Process (BP), cellular component (CC) and Molecular Function (MF). Figure showing processes with at least 20 genes involving at least 20% of total genes with enrichment p-value < 0.05.

### Integrated genome architecture of *P. cynomolgi* Berok

The Circos plot presented (Fig. 4A) provides an integrated, genome-wide view of the Berok strain genome assembly. The four concentric tracks (outer to inner) represent: (i) GC percentage (histogram), showing the sharp drop in GC content at the chromosome ends and in the SLE regions; (ii) gene annotations coloured by chromosome; (iii) stage-specific gene expression at 0, 12, 30, and 48 hours (standardized expression in red for high and blue for low), confirming that both core and SLE regions harbour transcriptionally active genes across the erythrocytic cycle; and (iv) alignment blocks against *P. cynomolgi* M (in green), highlighting the large Berok- specific (non-aligning) segments on chromosomes 4, 8, and 13. These tracks show that the SLEs, despite their extreme base composition and low gene density, are not inert heterochromatic deserts, instead they contain genes that are actively transcribed in a stage-specific manner and follow the just-in-time cascade observed across the core genome.

**Figure 4.**
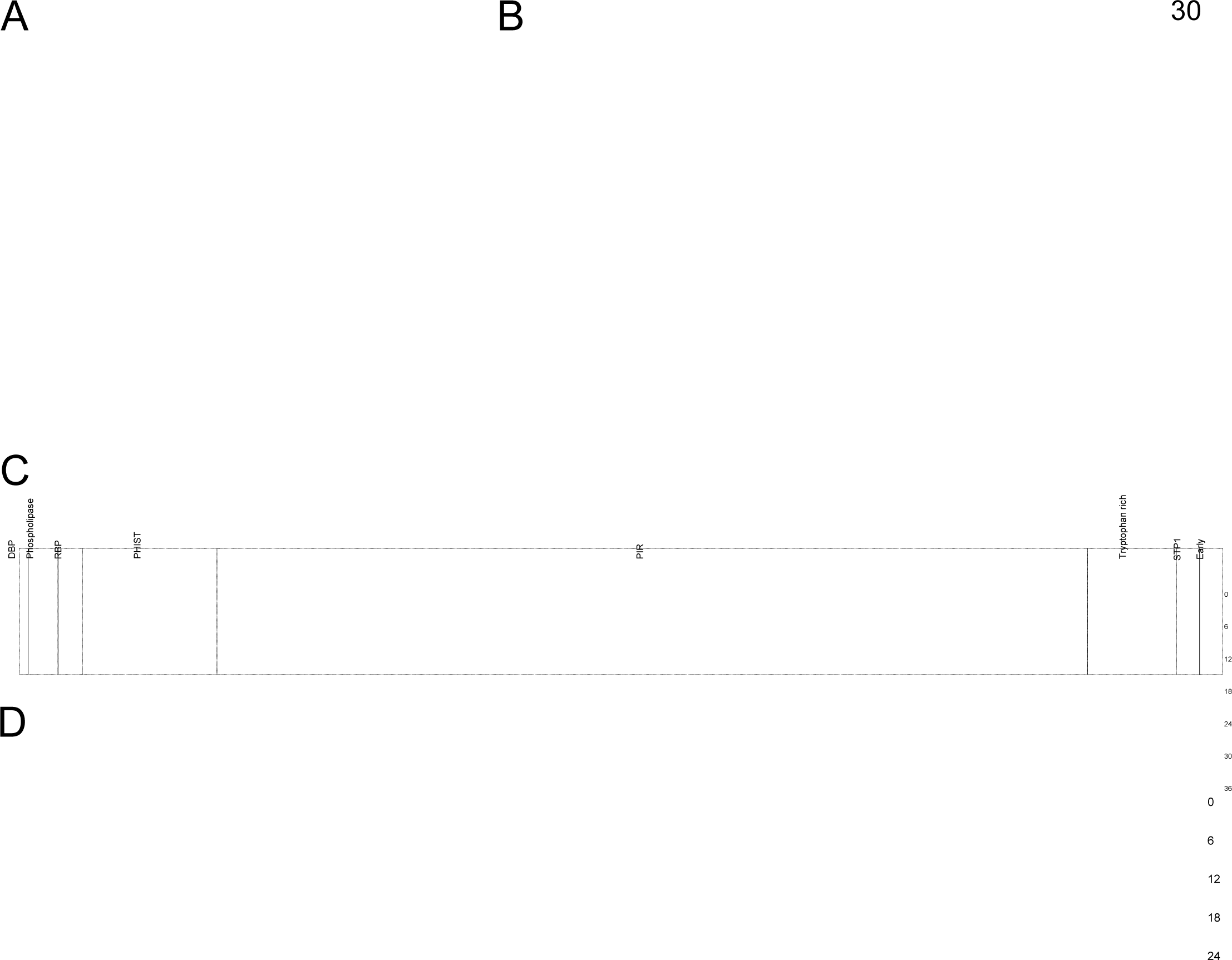
Integrated genome architecture of *P. cynomolgi* Berok. (A) The four concentric rings from outer to inner represent (i) GC percentage (histogram), (ii) gene annotations (colored by chromosome), (iii) gene expression at 0, 12, 30 and 48 (standardized expression in red and blue for high and low expression respectively) and (iv) alignments against *P. cynomolgi* M (in green). (B) Genes from the various invasion gene families (tryptophan-rich, STP1, RBP, PIR, PHIST, lysophospholipase, early invasion genes and DBP) depicted as dots on the *P. cynomolgi* Berok genome with the number of genes in each family on the table to the right showing the number and percentages. (C) Temporal expression of various invasion genes for IDC time- points. (D) Temporal expression of SLE genes for IDC time-points.

### Invasion gene families are enriched in subtelomeric regions and show coordinated stage- specific expression

Genome-wide mapping of invasion gene families including tryptophan-rich proteins, STP1, RBP, PIR, PHIST, lysophospholipase, early invasion genes, and DBP, reveals strong enrichment in subtelomeric and SLE regions (Fig. 4B; Supplementary Table 3). *PIR* genes are the most numerous (357 genes, 68.65%), followed by PHIST (68 genes, 13.08%). Several families accumulate preferentially at chromosome ends (e.g. RBP, tryptophan-rich), while others distribute more internally (e.g. PIR); chromosomes 4-7 and 10-12 harbour the greatest concentration of invasion gene loci. Temporal expression profiling across the IDC shows that these families display tightly coordinated, stage-specific transcriptional profiles (Fig. 4C), indicating that their subtelomeric enrichment reflects a preserved regulatory program rather than merely structural expansion. Taken together, these patterns point to a structured genomic organization that is consistent with family-specific expansion and diversification and suggest that *P. cynomolgi* Berok invests heavily in diversifying its surface repertoire, with a substantial proportion of exportome genes still of unknown function. The SLEs themselves are markedly enriched for these families: 178 of 274 SLE-encoded genes (64.96%) belong to an invasion gene family (Supplementary Fig. 7; Supplementary Table 4), with *PIR* genes again the most numerous (158 genes, representing 44.26% of all PIRs), followed by tryptophan-rich proteins (9 genes, 23.68%) and STP1 (5 genes, 50%). Of the remaining 96 SLE genes, 67 (69.80%) have no known function. Like the broader invasion gene complement, SLE genes show coordinated, stage-specific expression throughout the IDC (Fig. 4D), reinforcing the view that the SLEs constitute an actively regulated genomic compartment. This enrichment in multigene immune evasion families is consistent with the known biology of *Plasmodium* subtelomeres and supports the interpretation that the SLEs function as a structured reservoir for the diversification of host-interacting gene families. The high proportion of genes of unknown function within the SLEs (69.80% of non-invasion-family genes) further suggests that these compartments harbour biology that remains entirely unexplored.

### Phylogenomic divergence suggests a distinct Berok lineage

The core genome phylogeny (Fig. 5) mirrors the natural history of *Plasmodium*, organizing species by their long-term host associations. Avian parasites sit on a deeply separate branch from mammalian malaria, while rodent parasites cluster together, reflecting their shared evolutionary paths. Among primate parasites, *P. falciparum* groups closely with *P. reichenowi*, reinforcing their well-established shared ancestry. A second primate clade links *P. vivax* with the monkey malaria parasites *P. knowlesi* and *P. cynomolgi*, highlighting a close evolutionary continuum between human and non-human primate malaria species. Within *P. cynomolgi*, however, the Berok strain stands out as a deeply diverged and consistently supported lineage. This level of genome-wide separation suggests that Berok may represent more than simple strain variation and is consistent with the possibility of subspecies-level divergence.

**Figure 5.**
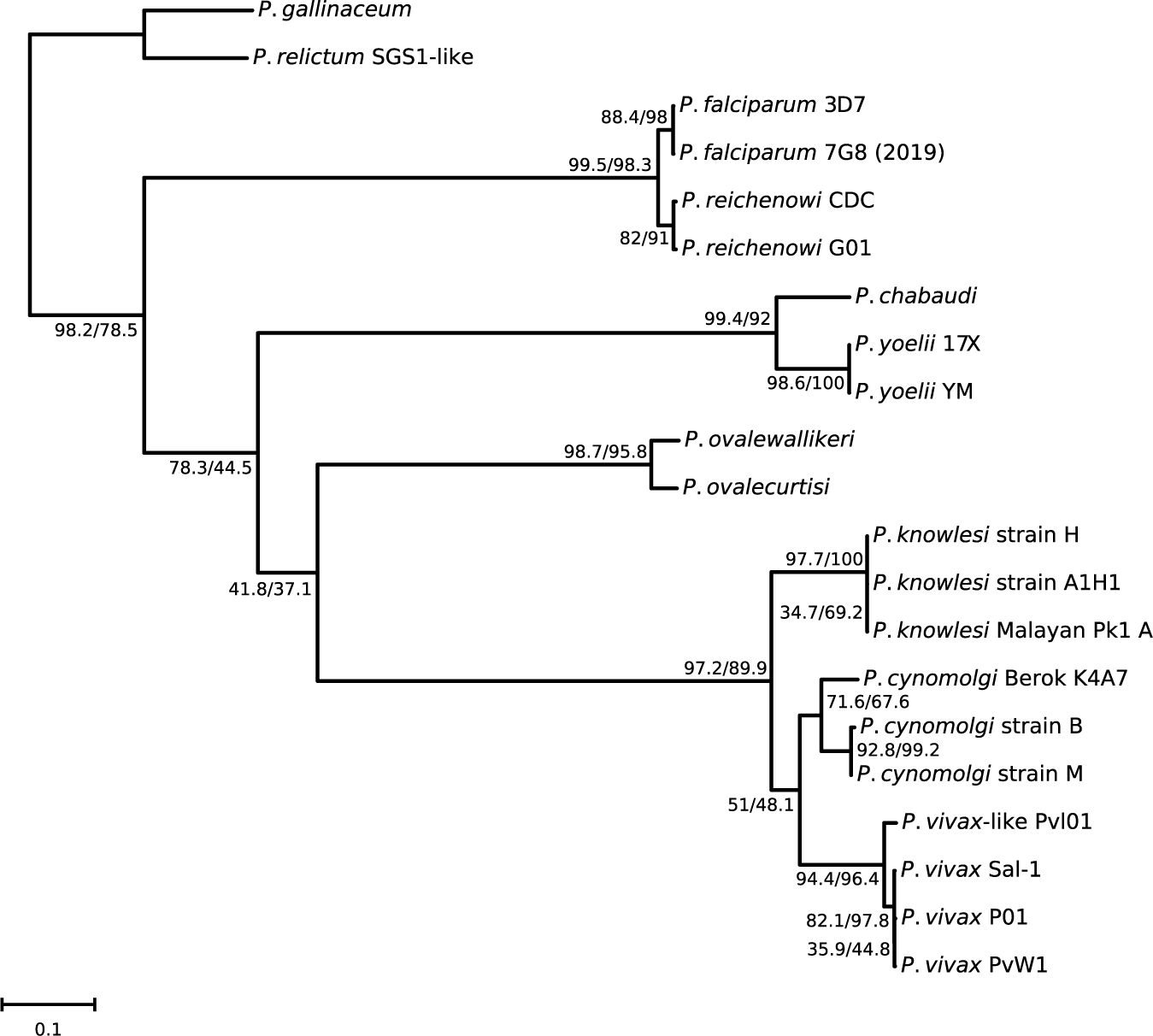
Phylogeny across selected *Plasmodium* species. Core genome phylogenetic tree of *Plasmodium* species and strains based on the concatenated amino acid sequences of 2603 single-copy orthologous genes. Branches are labelled with Gene Concordance Factor (gCF) and Site Concordance Factor (sCF), separated by a slash. Branch support was calculated via Ultrafast Bootstrap (UFBoot) with all values being 100% (not shown on tree). *Plasmodium gallinaceum* and *Plasmodium relictum* were used as the outgroup. The scale bar represents 0.1 substitutions per site.

Thus, pairwise comparisons of core single-copy orthologous genes were carried out. This shows that *P. cynomolgi* Berok and *P. cynomolgi* M differ by 16.7 amino acids per 100 aligned residues, a level of divergence comparable to that observed between other recognized *Plasmodium* species pairs such as *P. yoelii* and *P. chabaudi* (13.9) or *P. gallinaceum* and *P. relictum* (16.6). This level of divergence is substantially higher than that between the two *P. vivax* sequenced lines PvP01 and Sal-1 (0.8), but lower than the differences between the Berok strain and non- *cynomolgi* primate *Plasmodium* such as *P. vivax* (22.3) or *P. knowlesi* (23.3). This level of amino acid divergence would place *P. cynomolgi* Berok as a point intermediate between the levels of divergence within a species and that between species. These patterns illustrate how genome wide, quantitative metrics of divergence blur the simple dichotomy between “strain” and “species”, particularly in primate malaria parasites whose taxonomic boundaries were originally drawn on morphological or limited molecular grounds. These genome-wide comparisons show that the divergence between Berok and M is much greater than that usually seen between strains and therefore, must be considered when interpreting the taxonomic status of *P. cynomolgi* lineages. These data support the view that the Berok strain represents a markedly distinct lineage within *P. cynomolgi* populations. In this sense, Berok provides a concrete example of how geographically and host-associated lineages can accumulate substantial genomic divergence without yet being formally recognized as separate species, raising broader questions about species limits in zoonotic *Plasmodium* complexes (Supplementary Table 5). The same issue may apply to other zoonotic *Plasmodium* complexes in which deeply diverged lineages remain grouped under a single species name.

## Discussion

The availability of a complete and accurate genome sequence is of evident and substantial utility for investigations of all aspects of the life cycle and course of infection of a given *Plasmodium* species. It also provides invaluable data for future studies on the genetic diversity of the species populations, to resolve taxonomic status, and help derive an accurate phylogenetic positioning and history of *Plasmodium* parasites.

The chromosome-scale assembly of the *P. cynomolgi* Berok K4-A7 cloned line presented here represents a substantial advance over existing *P. cynomolgi* reference genomes, achieving near-complete chromosomal resolution with only 7 unplaced contigs representing a minute fraction of the whole genome (0,18%). This advance was made possible through the complementary use of sequencing and genome assembly strategies that made it possible to confidently place predominantly AT-rich contigs of many megabases that remained unplaced after routine sequencing/assembly approaches (Table 1). The two salient findings from this complete assembly are the substantially large genome size of the Berok strain (37 Mb as compared to 30,66 Mb for the M strain) principally from the 8 Mb resulting from our initial assembly, that were predominantly positioned as sub-telomeric expansions, and the significant divergence of the Berok strain genome from that of the M strain.

### The subtelomere-like expansions

The vast majority of the initially unplaced contigs totalling 8 Mb, were assigned to the subtelomeric regions of three chromosomes 4, 8 and 13. These SLEs are compositionally extreme (∼20% GC), gene-sparse, and enriched for multigene families coding for variable proteins exported to the infected erythrocyte membrane likely involved in host–parasite interactions such as the cynomolgi-interspersed repeat (CYIR) proteins, as well as methyltransferase pseudogenes (Supplementary Fig. 7). These large segments on chromosomes 4, 8, and 13 are absent from the M genomes. The k-mer composition analysis (Supplementary Fig. 6) and codon usage analysis (Supplementary Fig. 5) confirm that the SLE sequences are compositionally consistent with extreme AT-biased *Plasmodium* sequences and are not due to contamination, as validated by Kraken2 and Centrifuge analysis (Supplementary Fig. 3). The 93 *cyir* genes within the SLEs are of particular interest as the largest multigene family in *P. cynomolgi*. The expansion of *cyir* genes in the SLEs raises the possibility that the Berok lineage has a qualitatively different repertoire of immune evasion proteins compared to strains so-called B (M) and M, potentially relevant to its ability to establish chronic infections in macaque hosts and its zoonotic potential. Future functional studies using CRISPR-Cas9- mediated deletion of individual SLE-CYIR genes, coupled with *in vitro* adhesion and immune evasion assays, would be valuable for testing this hypothesis.

Beyond the CYIR family, the SLEs are broadly enriched for invasion gene families (Fig. 4B): 178 of 274 SLE genes (64.96%) belong to one of these families, with PIR genes being the most numerous (158 genes, 44.26% of all PIRs), followed by tryptophan-rich proteins and STP1. Critically, these genes are not transcriptionally silent. SLE-encoded genes display coordinated, stage-specific expression throughout the IDC that closely mirrors the temporal profiles of the broader invasion gene repertoire (Fig. 4C, 4D), and a subset is specifically upregulated in ring, trophozoite, and schizont stages. This places the SLEs firmly within the parasite’s active transcriptional program rather than outside it. The parallel between SLE gene expression and core genome invasion gene expression suggests a shared regulatory logic, raising the question of whether the same epigenetic machinery including H3K9me3 heterochromatin spreading and HP1 binding that governs *var*, *rifin*, and *stevor* expression in *P. falciparum*, also controls SLE gene regulation in *P. cynomolgi* Berok, a question now experimentally tractable given the available *in vitro* culture system.

Overall, *P. cynomolgi* Berok uses the same basic IDC script seen in other malaria parasites: a tightly phased cascade in which each gene is turned on when its product is needed. The smooth trajectory in expression space and the clean transcriptional waves argue that our time series captures this program with good temporal resolution and synchrony.

At the same time, the large number of phase-shifted orthologs shows that this script is not rigid. *P. vivax* and *P. cynomolgi* appear to run similar “scenes” in a different schedule, especially for genes involved in protein synthesis, degradation, and metabolism. Adjusting the timing of this core processes could help each species optimize resource use and IDC timing in its particular host environment, without changing the underlying protein-coding sequences. This set of 944 phase-shifted genes therefore provides a focused entry point for dissecting how regulatory changes in time, rather than in gene content, contribute to species-specific IDC biology and may influence traits such as stress responses and drug sensitivity.

A key finding is that SLE-encoded genes are transcriptionally active: lifecycle RNA-seq data confirm stage-specific expression of SLE-embedded genes during the erythrocytic cycle, with a subset upregulated in ring, trophozoite, and schizont stages (Fig. 3, Fig. 4A). This is consistent with findings in *P. falciparum*, where subtelomeric genes, including the *var*, *rifin*, and *stevor* families, display tightly regulated stage-specific expression despite a repetitive genomic context reinforcing the view that transcriptional activity in AT-rich subtelomeric compartments is a conserved feature of *Plasmodium* biology.

Resolving extreme subtelomeric sequences at chromosome scale is not a technical improvement, it is what makes their gene content, regulatory activity, and evolutionary dynamics interpretable in the first place. In *P. cynomolgi* Berok, anchoring the SLEs exposes an 8 Mb compartment that expands the genome’s host-interaction repertoire, harbours transcriptionally active multigene families, and integrates into the three-dimensional chromatin landscape of the erythrocytic cycle. This principle almost certainly applies beyond *P. cynomolgi*: in any *Plasmodium* lineage where subtelomeric regions remain fragmented or unplaced, the functional and evolutionary roles of these domains will be systematically underestimated.

### *P. cynomolgi* Berok, a lineage distinct from *P. cynomolgi* M

The phylogenomic analysis of the Berok and M strains clearly demonstrate that the two belong to distinct lineages. This was only suggested when the core genomes were analysed phylogenetically, which was re-enforced when divergence at the amino acid level was further analysed. The level of divergence far exceeds that between *P. falciparum* or *P. vivax* lineages collected in geographically distant locations, and approaches that between *P. cynomolgi* and *P. vivax*.

Historically four sub-species have been recognised *P. cynomolgi*^30^. *P. c. cynomolgi* is represented by the M strain that was initially isolated in the early 1930’s from an *M. fascicularis* imported to India from Singapore, *P. c. cyclopis* was recognized in 1960 as a subspecies that was described in the early 1942 from a *M. cyclopis* (Formosan rock macaque) from Taiwan, *P. c. bastianellii* was described in 1959 from an *M. fascicularis* from Peninsular Malaysia, and *P. c. ceylonensis* in 1965 from *M. sinica* (Toque macaque) and *Semnopithecus entellus* (Northern plain grey langur) from Sri Lanka. This classification was made by British parasitologists on morphological grounds, including those of the hepatic and sporogonic stages, though their colleagues in the United States consider that these merely represent distinct strains of *P. cynomolgi* ^31^. Thus, “*P. cynomolgi* B strain” is often used when referring to *P. c. bastianellii*, and “Ceylon Strain” for *P. c. ceylonensis*. Over the years numerous other *P. cynomolgi* isolates were collected throughout Southeast Asia^21^, though the two most often used experimentally are *P. c. bastianelli* and the M strain.

The notion that *P. cynomolgi* comprises distinct subpopulations was first provided by serological analysis targeting the circumsporozoite protein (CSP). Thereby, panel of monoclonal antibodies raised against five of the strains and tested against 11 strains tested revealed that these could be ascribed to at least five groups^32^, interestingly the Berok strain was the only one that cross-reacted with a monoclonal antibody raised against the *P. vivax* CSP. The suggestion that *P. cynomolgi* is a complex comprising multiple distinct lineages was reinforced when the sequences of the *csp* genes of six strains were found to have distinct repeat elements coding for the central repeat region of their respective CSPs^33^. Although some *P. cynomolgi* genes were sequenced over the next years, the first studies on the diversity of *P. cynomolgi* was published 20 years later and focused the merozoite surface antigen 1 (*msp1*) gene from 11 strains that included three subspecies^34, 35^, and subsequently included the mitochondrial genome^36^ and the MSP8 and MSP10 genes^37^. In these studies, the phylogenetic analyses showed that the Berok and Gombak strains invariable clustered together and separately from the other strains analysed.

Recently surveys of the parasites in naturally infected macaques, particularly in Malaysia, provided the opportunity to study the genetic diversity of *P. cynomolgi*. Sequences from four genes were investigated, that coding for the apical membrane antigen 1 (AMA1)^3^, the apicoplast caseinolytic protease M (ClpM) and the mitochondrial genome^38^, and Region II of the Duffy binding proteins 2 and 1 (DBP2 and DBP2)^39, 40^. In all case phylogenetic analyses showed that Berok as a separate lineage from other known strains. Moreover, these analyses also revealed that Berok-type parasites are circulating in nature, and one of the studies suggested that another a novel *P. cynomolgi* lineage, as well as others from other *Plasmodium* species, might be present in macaques from Borneo^3^. Given the consistency with which the Berok and Gombak strains cluster in a lineage distinct from that of the other strains and subspecies, we propose that these two strains represent members of a taxonomically distinct lineage within *P. cynomolgi* meriting subspecies status, we propose the designation of *Plasmodium cynomolgi beroki* n. subsp. (pending formal nomenclatural revision with appropriate material designation). The genetic distance between the Berok and M strains and the fact that the Berok/Gombak lineages are sympatric with other *P. cynomolgi* lineages is akin to the situation with the *P. ovale* populations that were found to comprise two distinct species, *P. ovalecurtisi* and *P. ovalewallikeri*^41^. Analyses of whole genome sequences from the other lineages combined with a lack of evidence of hybridisation between them might lead to elevating *P. c. beroki* to the species level. This has important implications for interpreting cross-strain comparisons and for defining the host range and epidemiological relevance of different *P. cynomolgi* lineages.

### P. cynomolgi beroki as a model for P. vivax

A listing of all the potential genes, pseudogenes, and other genome structures from the near-complete genome of *P. c. beroki*, provides a potential insight into many of the biological processes over the life cycle of this parasite. Methodologically, our approach consisting of anchoring a near complete genome, quantifying genome wide divergence across thousands of orthologs, and integrating historical single gene and mitochondrial data, provides a reusable framework for revisiting other *Plasmodium* complexes where species limits remain debated. The combination of a near-complete genome assembly with genome-wide phylogenetic comparisons should also be useful for examining other *Plasmodium* complexes in which species limits remain uncertain. This is all the more useful because this subspecies, as well as the various *P. cynomolgi* lineages, are morphologically indistinguishable from their genetically closest relative *P. vivax*, and display similar biological activities, including hypnozoite formation. However, investigations of these parasites have been limited to collected blood samples for *P. vivax*, a parasite that is not amenable to *in vitro* cultivation, and materials and observations from infected Rhesus monkeys. The availability of the *P. c. beroki* K4 line and its cloned line K4A7 from continuous *in vitro* culture can now facilitate leveraging the genome sequence to address a number of previously intractable functional investigations.

First, improvements in *in vitro* cultivation protocols reduce the need for macaque serum since robust growth is obtained using foetal bovine serum or normal horse serum^16^, and an eventual adaptation to multiplication in human red blood cells would be a major enhancement. The K4A7 cloned line has also been shown to be amenable to genetic modification via CRISPR-Cas 9^17^ in a study that further demonstrated its proven utility for drug susceptibility investigations^42^. This opens the way to conduct detailed and tractable investigations on the biology of the erythrocytic stages, such as the function of the caveola-vesicle complex or Schüffner dots^43^, and the mechanism of red blood cell invasion selectivity. Genes coding for orthologous and paralogous proteins implicated in red blood invasion have been identified in the genomes of *P. vivax* and *P. cynomolgi*. Their structures and numbers vary between the species and between the strain’s studies, leading to tantalising speculations as to the actual functional significance of such differences. The functional role of these proteins can now be directly addressed in experimental studies akin to the elegant investigations using the less genetically related *P. knowlesi*^44, 45, 46, 47^. Hypnozoites are of particular interest to the malaria community, as the relapses they cause are a major hurdle in efforts to control *P. vivax*. *P. cynomolgi* has served in the seminal discovery of this dormant hepatic form and has long been the model used to identify drugs to inhibit it. The development of *P. cynomolgi* hepatic *in vitro* cultures revealed the presence of uninuclear forms that behave similarly to hypnozoites^5^ and that are used to study hypnozoite biology^48^, or to identify the genes that might be implicated^49, 50, 51, 52^. Incrimination of a particular gene will, however, require *in vivo* confirmation through genetically modified parasites that could be generated in *P. c. beroki*. It must be noted that this parasite, as are other species that infect macaques, remain the only *Plasmodium* species that can be experimentally investigated throughout the course of natural infections in their natural host *M. fascicularis*.

In conclusion, the complete genome of *P. c. beroki* and the ability to cultivate it *in vitro* makes this parasite line highly suitable as an experimental proxy for *P. vivax*, with a heightened potential to help elaborate efficacious control measures.

## Materials and Methods

### Ethics and animal welfare

*Macaca fascicularis* monkeys serving as blood donors were maintained in a malaria-free facility at Singhealth Experimental Medicine Center, Singapore. All procedures were approved by Singhealth IACUC (2025/SHS/2002).

### Parasite culture and clonal selection

*P. cynomolgi* Berok K4 continuous culture was maintained as previously described. Clone K4- A7 was isolated by limiting dilution (0.5% parasitaemia, 1.25% haematocrit; 10-fold serial dilution across 96-well plates) with biweekly RBC supplementation and identified at day 55 by flow cytometry (dihydroethidium/Hoechst staining) and Giemsa confirmation.

### Genomic DNA extraction

Mixed-stage cultures (∼2 × 10⁹ parasites) were lysed with 0.15% saponin/0.1% BSA in PBS. DNA was isolated using Blood and Cell Culture DNA Mini extraction kit (Qiagen) and quantified by Quant-iT PicoGreen assay (Invitrogen). Quality was assessed using LabChip GX (PerkinElmer).

### Oxford Nanopore sequencing

Libraries were constructed from 2.5 μg gDNA using the 1D² adapter ligation kit (SQK-LSK-308, ONT) and sequenced on GridION (R9.5.1 chemistry, FLO-MIN 107) for 48 hours. Base calling was performed with Guppy 1.5.1-1; only reads from the first 24 hours (higher quality) exceeding 1 kb were retained.

### Illumina sequencing

PCR-free libraries were prepared using TruSeq DNA PCR-Free Library Prep Kit (Illumina) from 2 μg gDNA, fragmented and size-selected (550 bp) with Ampure beads (Beckman Coulter). Libraries were sequenced on MiSeq generating 2 × 251 bp paired-end reads.

### Hi-C library preparation

Hi-C data were generated using the Phase Genomics Proximo Hi-C 4.0 Kit. Intact cells were crosslinked with formaldehyde, HMW DNA was extracted (Qiagen), digested with DPNII/DdeI/HinfI/MseI, and proximity ligated with biotinylated nucleotides. Chimeric molecules were captured with streptavidin beads and sequenced on Illumina NovaSeq generating 81.5 million PE150 read pairs.

### Hybrid assembly and scaffolding

Nanopore reads >1 kb were assembled *de novo* with CANU (version 2.1). Contigs were polished sequentially: Racon (version 1.4.13) (with minimap2-aligned raw reads), Nanopolish (version 0.13.2) (using original fast5 signal data), and Racon again (with BWA-aligned Illumina reads). Hi-C reads were aligned to contigs using BWA-MEM (−5SP -t 8) (version 0.7.17-r1198-dirty), PCR duplicates flagged with SAMBLASTER (version 0.1.24), and non-primary alignments filtered with samtools (-F 2304) (version 8.22). Chromosome-scale scaffolding was performed using Phase Genomics’ Proximo platform and manually corrected with Juicebox (version 2.15).

### RNA-seq and transcriptomic analysis

Synchronized parasites (∼90% schizonts via MACS enrichment) were added to naïve *M. fascicularis* RBCs at ∼1% parasitaemia. Samples were harvested every 6 hours from 0 to 54 hours post-setup for RNA-seq analysis with RPKM quantification.

RNA-Seq read sequences were obtained in FASTQ format. The quality of the FASTQ files were checked using FASTQC. The reads were aligned to the assembled genome using minimap2 (version 2.30) and gene counts computed using featureCounts (version 2.0.3) from the subread package using the genome annotations generated.

### Core genome phylogenetic tree

OrthoFinder^53, 54^ version 3.1.4 was used to obtain single-copy orthologous genes (SCOGs) from amino acid sequences between *Plasmodium cynomolgi* Berok K4A7 and 20 *Plasmodium* species and strains from PlasmoDB^55^ (Supplementary Table 5 and 6). MAFFT^56^ version 7.526 with automatic algorithm selection was used to generate alignments for each of the 2606 SCOGs obtained. TrimAl^57^ version 1.5.1 was used for trimming on each alignment using the “gappyout” mode. Three trimmed alignments which comprised solely of identical sequences were excluded. The remaining alignments were concatenated into a supermatrix with a corresponding partition file using AMAS^58^ version 1.0. IQTREE3^59^ version 3.1.1 was then used to derive a phylogenetic tree from the supermatrix with 1000 Ultrafast Bootstrap^60^ replicates. IQTREE3 was also used to compute gene trees from the individual alignments. A seed of 0 and the MFP+MERGE option to obtain the best substitution model for each partition was used for IQTREE3 runs^61, 62, 63^. IQTREE3 was then used to compute for each branch the Gene Concordance Factor (gCF), as well as Site Concordance Factor (sCF) with 100 replicates (--scfl 100)^64^. ETE 3^65^ was used to reroot the tree using *Plasmodium gallinaceum* and *Plasmodium relictum* as an outgroup. BioPython^66^ version 1.86 was used for tree plotting.

### Calculation of pairwise percentage amino acid identity of SCOGs

BioPython version 1.86^66^ was used to extract and compare amino acid sequences from all 2606 SCOGs identified by OrthoFinder for specified pairwise comparisons of *Plasmodium* species or strains. For each pairwise comparison, the two corresponding sequences for every SCOG were aligned using the Needleman-Wunsch algorithm with the BLOSUM62 substitution matrix (cysteines replacing selenocysteines for BLOSUM62 compatibility). Any resulting overhangs were trimmed. Percentage identity for each SCOG was computed via dividing the number of matches by the trimmed alignment length, then multiplying the value by 100. To obtain summary statistics, median percentage identity of all SCOGs was computed, as well as the 1^st^ and 3^rd^ quartiles.

### BUSCO quality assessments

BUSCO^67, 68^ version 6.0.0 was used with plasmodium_odb12 (2025-07-01) as the lineage dataset and Metaeuk^69^ as the gene predictor. The *Plasmodium cynomolgi* B and M strain genomes were retrieved from https://plasmodb.org/common/downloads/Current_Release/PcynomolgiB/fasta/data/PlasmoDB-68_PcynomolgiB_Genome.fasta and https://plasmodb.org/common/downloads/Current_Release/PcynomolgiM/fasta/data/PlasmoDB-68_PcynomolgiM_Genome.fasta respectively on 31 March 2026.

### Data availability

Sequence data will be deposited in INSDC (GenBank/ENA/DDBJ) and RNA-seq data in GEO/SRA.

## Acknowledgments

The work was supported by in part by the Lee Kong Chian School of Medicine Start-Up Grant (award number: 025277-00011) and the A*STAR Infectious Diseases Labs core research grants by the Biomedical Research Council (BMRC) of Agency for Science, Technology and Research (A*STAR) and TF IPC Ltd grant titled “Temasek Foundation Infectious Diseases Programme for Surveillance and Diseases X Resilience.” awarded to P. B. Z.B. is partly supported by a Singaporean Ministry of Education Grant (grant number MOE2019-T3-1-007 to ZB) P.C. and B.R. are supported by the University of Otago, New Zealand (Marsden University of Otago, Deans Bequest Grant). A.C.Y.C. was partly supported by the National Medical Research Council (NMRC), Singapore, Open Fund Young Individual Research Grant (OF-YIRG), under grant IGMS-MOH-001727-00. LR was supported by the Singapore Ministry of Health Start-Up Grant [#SUG #022388-00001), the Ministry of health Tier 2 (#T2EP30125_0033) and by the National Medical Research Council (#OFIRG24jul-0055). IDMIT infrastructure (G.S.) is supported by the French “Programme d’Investissements d’Avenir” (PIA) under grant ANR-11-INBS-0008. G.S. is supported by Agence Nationale de la Recherche (ANR-17-CE13-0025-01). The SIgN Immunomonitoring platform is supported by BMRC IAF 311006 and BMRC transition funds #H16/99/b0/011.

## Author contributions

A.C.Y.C., G.S., and P.B. conceived the study. A.C.Y.C., P.C., S.K.G., V.C., K.S.S., S.X.T., R.S., and B.R., conducted the experiments. V.N., E.J.K.L., S.N., A.P., Z.B., and B.T.K.L., performed bioinformatic analyses. D.G. performed Hi-C library preparation and scaffolding. K.S.W.T, P.P, and L.R, provided structural support and critical analysis. A.C.Y.C., G.S., Z.B., and P.B. wrote the manuscript. All authors reviewed and edited the manuscript.

The authors declare no competing interests.

