## Supplementary Information for "A chromosome-scale Plasmodium cynomolgi Berok genome reveals a distinct subtelomeric architecture and a highly diverged primate malaria lineage"

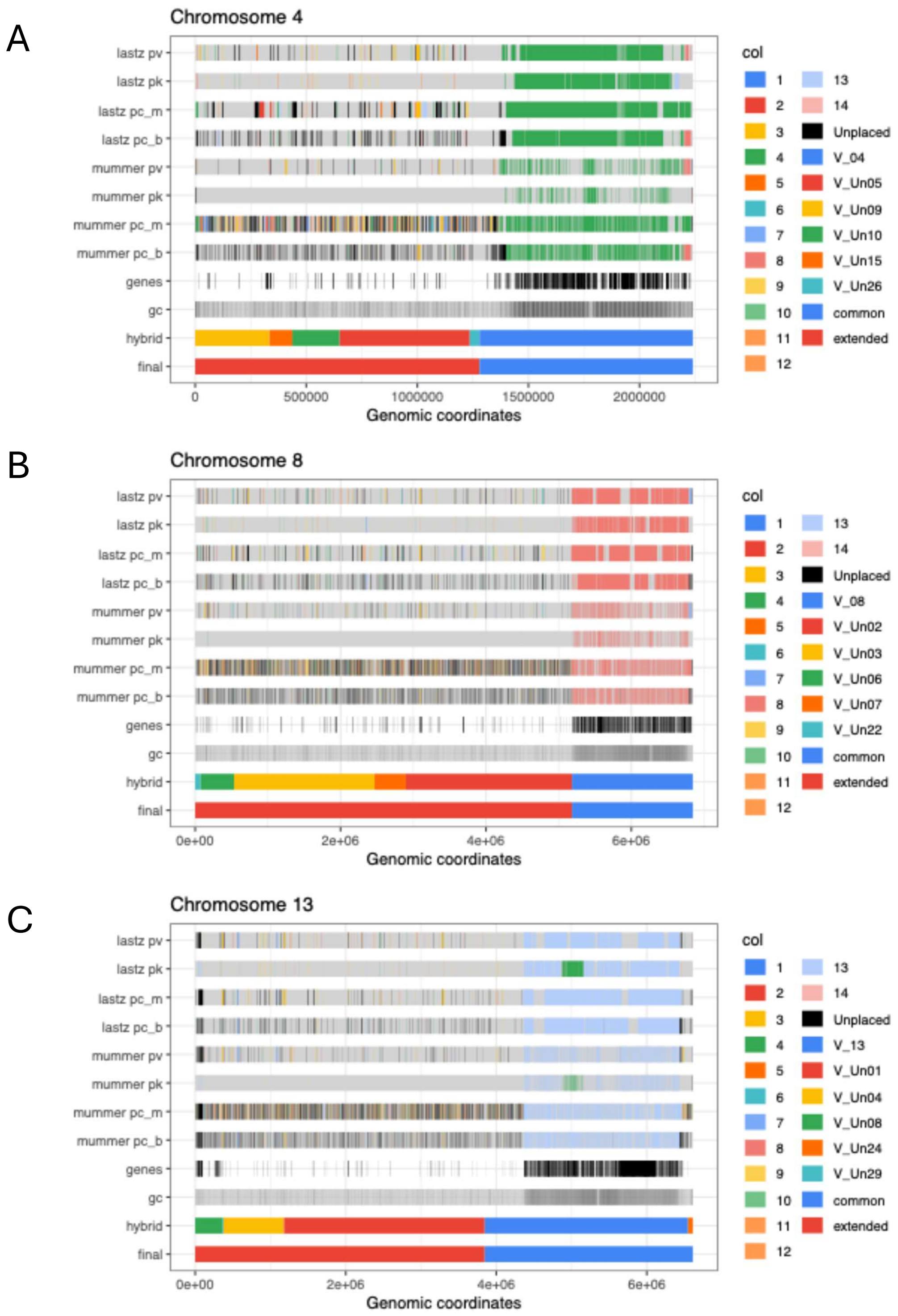


**Supplementary Figure 1. Plots showing the common and expanded regions in chromosomes 4 (A), 8 (B) and 13 (C).** The LASTZ and MUMMER aligned regions against the various *Plasmodium* species are colored by the chromosome number showing concordant assignment. The location of the genes are indicated in the row labelled genes while the GC content is indicated in the row labelled gc. The original contigs assembled using a hybrid approach prior to final assembly by Hi-C data is labelled as hybrid and the final annotation of expanded (new in Berok) and common (common to all analyzed *Plasmodium* species) is provided as the last row.

**Supplementary Table 1**: **Table showing the common regions as well as the expanded regions for each of the chromosomes.** The percentage of the total chromosome length (% of chr), GC content (GC %), number of genes and number of genes per 1Mbp for each region is shown. The table shows that the expanded regions are AT rich and that there is a low density of genes.

| Chr | Region | Start | End | Chr length | % of chr | GC % | No of genes | Number of gene per Mbp |
|---|---|---|---|---|---|---|---|---|
| 1 | common | 1 | 1,299,006 | 1,299,006 | 100.00 | 36.85% | 215 | 165.51 |
| 2 | common | 1 | 1,399,388 | 1,399,388 | 100.00 | 33.51% | 220 | 157.21 |
| 3 | common | 1 | 1,411,064 | 1,411,064 | 100.00 | 37.62% | 270 | 191.34 |
| 4 | expanded | 1 | 1,279,827 | 2,239,968 | 57.14 | 23.76% | 45 | 35.16 |
| 4 | common | 1,279,828 | 2,239,968 | 2,239,968 | 42.86 | 40.15% | 225 | 234.34 |
| 5 | common | 1 | 1,369,445 | 1,369,445 | 100.00 | 40.80% | 327 | 238.78 |
| 6 | common | 1 | 1,174,169 | 1,174,169 | 100.00 | 41.57% | 257 | 218.88 |
| 7 | common | 1 | 1,617,441 | 1,617,441 | 100.00 | 41.37% | 389 | 240.50 |
| 8 | expanded | 1 | 5,183,250 | 6,846,290 | 75.71 | 22.69% | 136 | 26.24 |
| 8 | common | 5,183,251 | 6,846,290 | 6,846,290 | 24.29 | 42.28% | 430 | 258.56 |
| 9 | expanded | 1 | 2,215,980 | 2,215,980 | 100.00 | 40.15% | 536 | 241.88 |
| 10 | common | 1 | 1,405,730 | 1,405,730 | 100.00 | 41.89% | 361 | 256.81 |
| 11 | common | 1 | 2,611,300 | 2,611,300 | 100.00 | 38.19% | 558 | 213.69 |
| 12 | common | 1 | 3,640,341 | 3,640,341 | 100.00 | 38.35% | 848 | 232.95 |
| 13 | expanded | 1 | 3,847,215 | 6,610,846 | 58.20 | 22.08% | 93 | 24.17 |
| 13 | common | 3,847,216 | 6,610,846 | 6,610,846 | 41.80 | 37.14% | 541 | 195.76 |
| 14 | common | 1 | 3,057,228 | 3,057,228 | 100.00 | 40.25% | 796 | 260.37 |

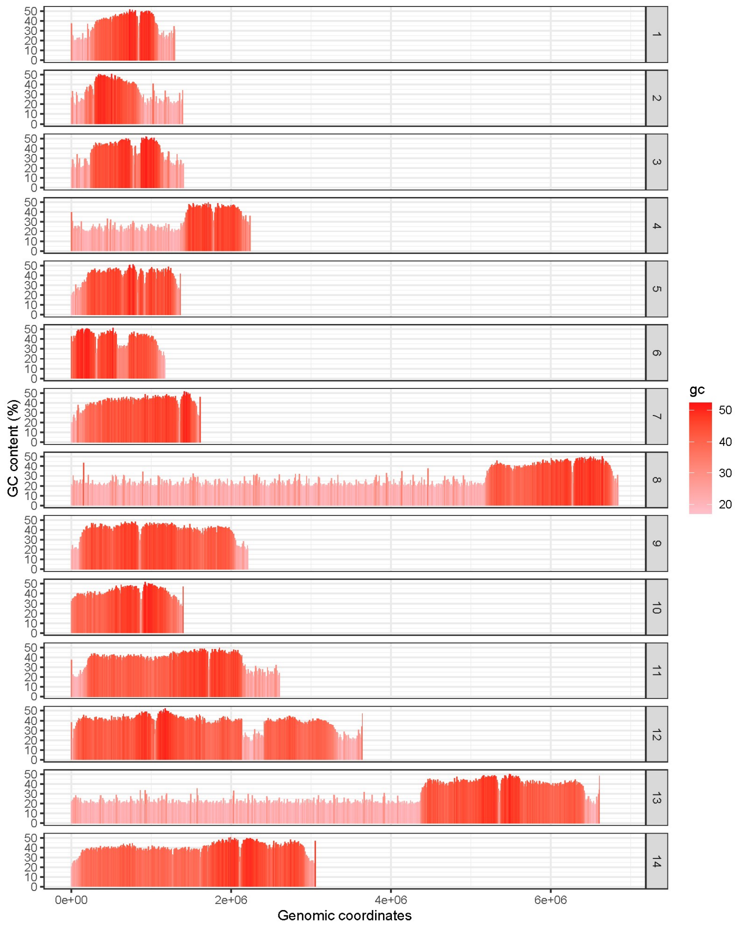


**Supplementary Figure 2. GC content of P . *cynomolgi* Berok.** GC content across all chromosomes. All chromosome ends and SLE regions (chromosomes 4, 8 and 13) are more AT rich (%).

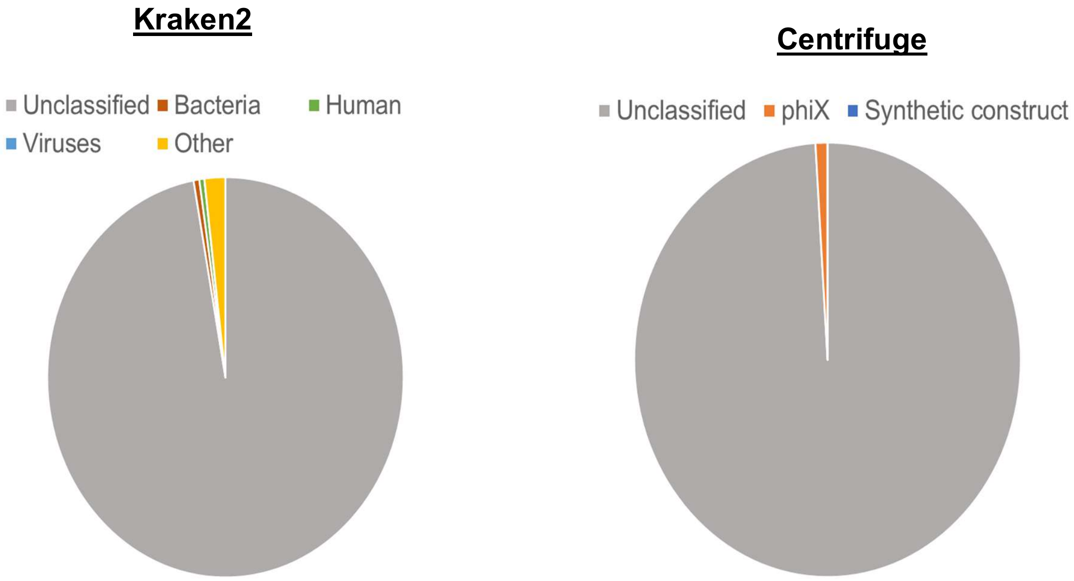


**Supplementary Figure 3.** Analysis was conducted to ensure that extra chromosomal DNA is not due to contamination.

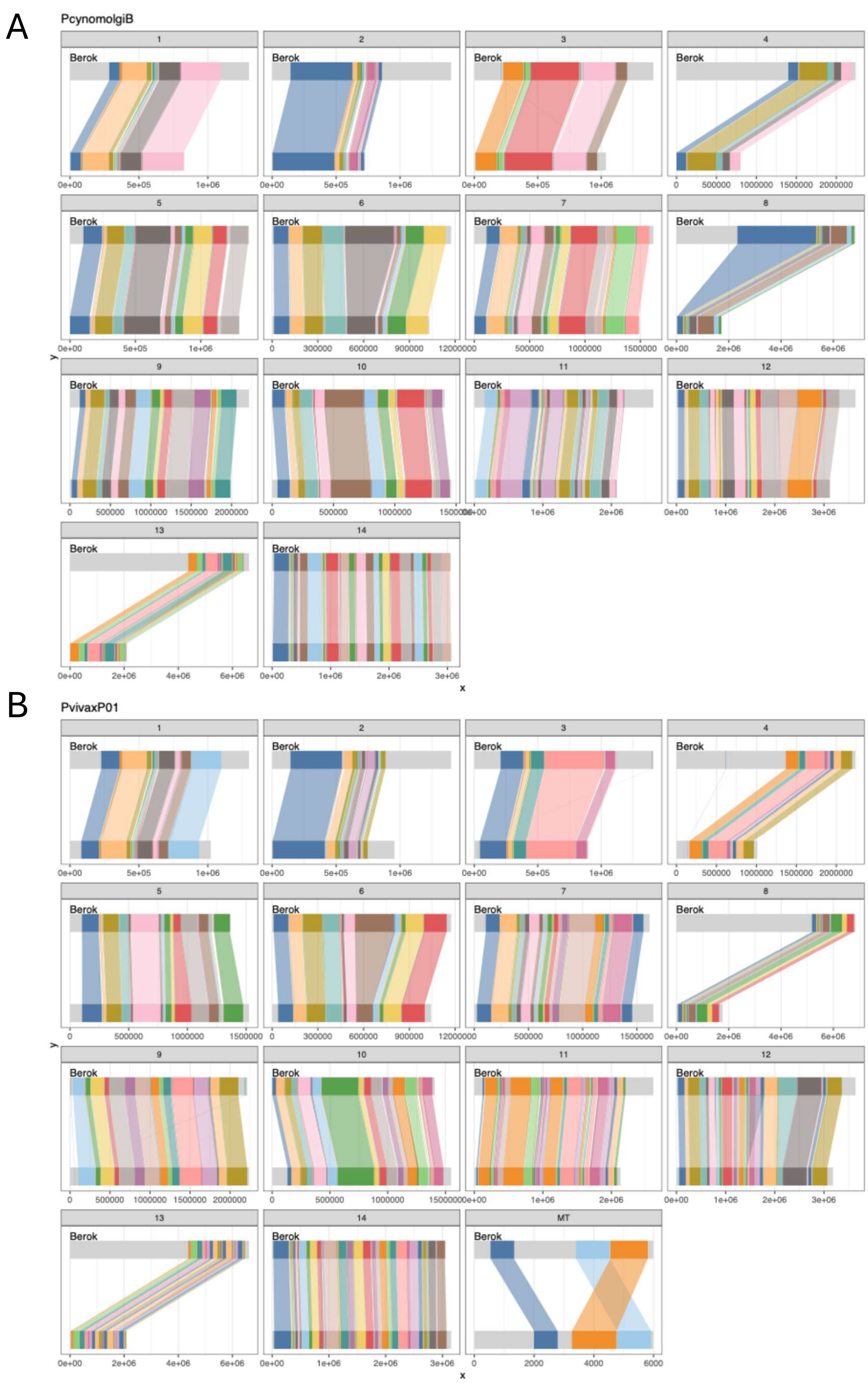


**Supplementary Figure 4.** Plots showing gene level synteny of (A) *P. cynomolgi* strain B and (B) *P. vivax* P01.

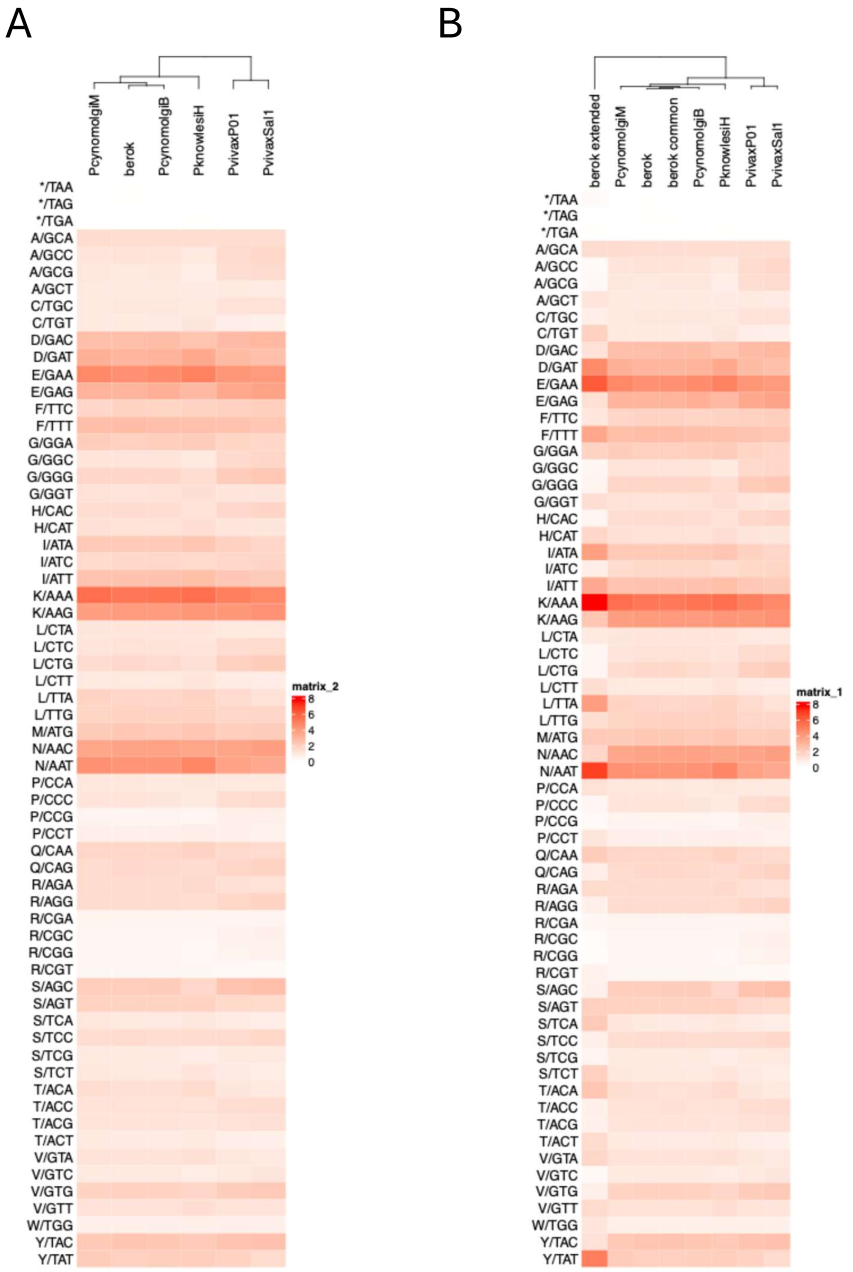


**Supplementary Figure 5. Heat map showing the codon usage across the various *Plasmodium* species.** (A) Heat map showing just the combined Berok codon usage showing the similarity between all *Plasmodium* species. (B) Heat map colored by the codon usage percentage (against total codons) with the Berok strain data shown as combined for the SLE and common regions (Berok) as well as individually (Berok SLE and Berok common).

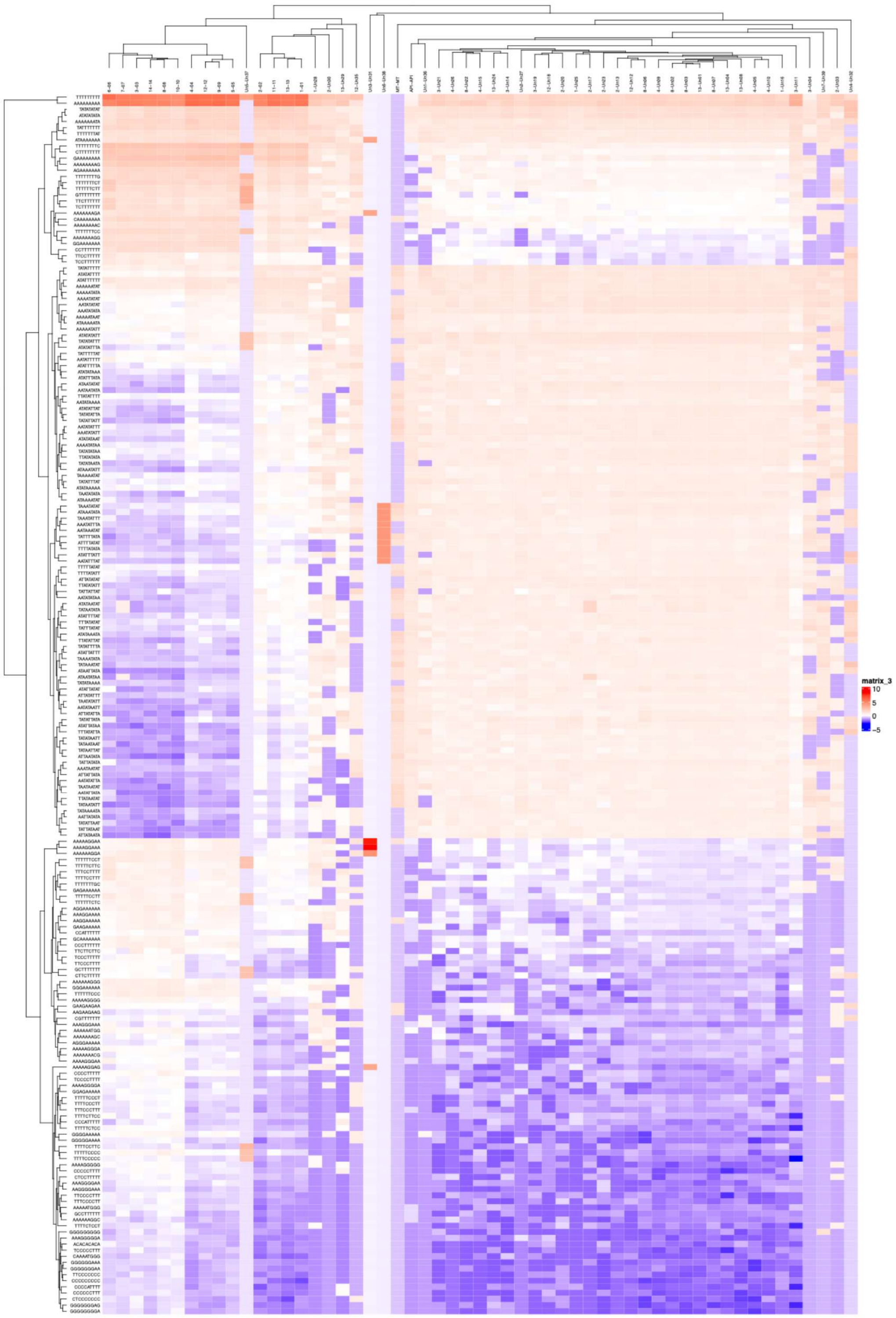
matrix_3
10
5
0
−5

**Supplementary Figure 6. Heat map showing the enrichment of specific 9mers in either the extended regions or common regions with other *Plasmodium* species.** The sequence of the 9-mers are indicated in the row names. The cells show the percentage of the 9mers for each of the chromosome/contigs with Z-score normalization.

**Supplementary Table 2:** The raw and normalized data of transcriptional profile of the corresponding temporal state of *P. cynomolgi* IDC. (Excel File)

**Supplementary Table 3**: Table of invasion genes showing the genomic location of each gene. A summary of the gene families is also provided. (Excel File)

**Supplementary Table 4:** Table of SLE genes showing the genomic location of each gene. (Excel File)

**Supplementary Table 5:** Summary statistics for percentage amino acid identity of 2606 Single Copy Orthologous Genes (SCOGs) between pairs of *Plasmodium* species or strains.

| Pairwise comparison | Median percenta ge amino acid identity | 1st quartile | 3rd quartile |
|---|---|---|---|
| ***P. cynomolgi* Berok K4A7 vs. *P. cynomolgi* strain M** | 91 | 82.5 | 96.4 |
| ***P. cynomolgi* Berok K4A7 vs. *P. cynomolgi* strain B** | 85.5 | 69.8 | 94.2 |
| ***P. cynomolgi* strain M vs. *P. cynomolgi* strain B** | 97.2 | 86.7 | 100 |
| ***P. cynomolgi* Berok K4A7 vs. *P. vivax* P01** | 83.3 | 72.3 | 92.1 |
| ***P. cynomolgi* Berok K4A7 vs. *P. vivax* Sal-1** | 83.3 | 72.6 | 92 |
| ***P. cynomolgi* Berok K4A7 vs. *P. vivax* PvW1** | 83.2 | 72.4 | 92 |
| ***P. cynomolgi* Berok K4A7 vs. *P. vivax-like* Pvl01** | 82 | 70.1 | 91.4 |
| ***P. cynomolgi* Berok K4A7 vs. *P. knowlesi* strain H** | 81.8 | 71.1 | 90.3 |
| ***P. cynomolgi* Berok K4A7 vs. *P. knowlesi* strain A1H1** | 81.8 | 71 | 90.2 |
| ***P. cynomolgi* Berok K4A7 vs. *P. knowlesi* Malayan Pk1 A** | 81.8 | 71 | 90.3 |
| ***P. cynomolgi* Berok K4A7 vs. *P. ovalecurtisi*** | 57.3 | 44.6 | 73.1 |
| ***P. cynomolgi* Berok K4A7 vs. *P. ovalewallikeri*** | 56.9 | 44.3 | 72.9 |
| ***P. cynomolgi* Berok K4A7 vs. *P. yoelii* 17X** | 53.4 | 41.6 | 67.9 |
| ***P. cynomolgi* Berok K4A7 vs. *P. yoelii* YM** | 53.4 | 41.6 | 67.8 |
| ***P. cynomolgi* Berok K4A7 vs. *P. chabaudi*** | 53.7 | 41.9 | 68 |
| ***P. cynomolgi* Berok K4A7 vs. *P. reichenowi* CDC** | 52.7 | 39.8 | 68.4 |
| ***P. cynomolgi* Berok K4A7 vs. *P. reichenowi* G01** | 52.7 | 39.7 | 68.6 |
| ***P. cynomolgi* Berok K4A7 vs. *P. falciparum* 3D7** | 52.6 | 39.8 | 68.8 |
| ***P. cynomolgi* Berok K4A7 vs. *P. falciparum* 7G8 (2019)** | 52.6 | 39.8 | 68.8 |
| ***P. cynomolgi* Berok K4A7 vs. *P. gallinaceum*** | 55.4 | 42.8 | 70.8 |
| ***P. cynomolgi* Berok K4A7 vs. *P. relictum* SGS1-like** | 55.8 | 43.2 | 70.7 |
| ***P. cynomolgi* strain M vs. *P. vivax* P01** | 86.6 | 78.1 | 93.8 |
| ***P. cynomolgi* strain M vs. *P. vivax* Sal-1** | 86.5 | 78.2 | 93.8 |
| ***P. cynomolgi* strain M vs. *P. vivax* PvW1** | 86.5 | 78.2 | 93.8 |
| ***P. cynomolgi* strain M vs. *P. vivax-like* Pvl01** | 85.4 | 76.3 | 93.3 |
| ***P. cynomolgi* strain M vs. *P. knowlesi* strain H** | 84.6 | 76.2 | 92.3 |
| ***P. cynomolgi* strain M vs. *P. knowlesi* strain A1H1** | 84.5 | 76 | 92.2 |
| ***P. cynomolgi* strain M vs. *P. knowlesi* Malayan Pk1 A** | 84.5 | 76 | 92.2 |
| ***P. cynomolgi* strain M vs. *P. ovalecurtisi*** | 60.3 | 47.8 | 75.5 |
| ***P. cynomolgi* strain M vs. *P. ovalewallikeri*** | 59.9 | 47.5 | 75.2 |
| ***P. cynomolgi* strain M vs. *P. yoelii* 17X** | 56.1 | 44.5 | 70.6 |
| ***P. cynomolgi* strain M vs. *P. yoelii* YM** | 56 | 44.5 | 70.5 |
| ***P. cynomolgi* strain M vs. *P. chabaudi*** | 56.3 | 45 | 70.3 |
| ***P. cynomolgi* strain M vs. *P. reichenowi* CDC** | 55.5 | 43 | 71 |
| ***P. cynomolgi* strain M vs. *P. reichenowi* G01** | 55.6 | 42.9 | 71 |
| ***P. cynomolgi* strain M vs. *P. falciparum* 3D7** | 55.6 | 43 | 71.1 |
| ***P. cynomolgi* strain M vs. *P. falciparum* 7G8 (2019)** | 55.6 | 43.1 | 71.1 |
| ***P. cynomolgi* strain M vs. *P. gallinaceum*** | 58.3 | 46.3 | 72.6 |
| ***P. cynomolgi* strain M vs. *P. relictum* SGS1-like** | 58.9 | 46.6 | 72.6 |
| ***P. cynomolgi* strain B vs. *P. vivax* P01** | 80.5 | 67.8 | 90.8 |
| ***P. cynomolgi* strain B vs. *P. vivax* Sal-1** | 80.6 | 68 | 90.8 |
| ***P. cynomolgi* strain B vs. *P. vivax* PvW1** | 80.5 | 67.7 | 90.7 |
| ***P. cynomolgi* strain B vs. *P. vivax-like* Pvl01** | 79.3 | 65.9 | 90.2 |
| ***P. cynomolgi* strain B vs. *P. knowlesi* strain H** | 79.1 | 66.7 | 89.1 |
| ***P. cynomolgi* strain B vs. *P. knowlesi* strain A1H1** | 79 | 66.5 | 89 |
| ***P. cynomolgi* strain B vs. *P. knowlesi* Malayan Pk1 A** | 79.1 | 66.6 | 89.1 |
| ***P. cynomolgi* strain B vs. *P. ovalecurtisi*** | 56 | 43.3 | 72.2 |
| ***P. cynomolgi* strain B vs. *P. ovalewallikeri*** | 55.5 | 43.1 | 72.2 |
| ***P. cynomolgi* strain B vs. *P. yoelii* 17X** | 52.1 | 40.3 | 67.1 |
| ***P. cynomolgi* strain B vs. *P. yoelii* YM** | 52.1 | 40.3 | 67.1 |
| ***P. cynomolgi* strain B vs. *P. chabaudi*** | 52.4 | 40.8 | 67.4 |
| ***P. cynomolgi* strain B vs. *P. reichenowi* CDC** | 51.5 | 38.6 | 67.6 |
| ***P. cynomolgi* strain B vs. *P. reichenowi* G01** | 51.6 | 38.4 | 67.6 |
| ***P. cynomolgi* strain B vs. *P. falciparum* 3D7** | 51.6 | 38.5 | 67.9 |
| ***P. cynomolgi* strain B vs. *P. falciparum* 7G8 (2019)** | 51.5 | 38.5 | 67.9 |
| ***P. cynomolgi* strain B vs. *P. gallinaceum*** | 54.4 | 42 | 70.1 |
| ***P. cynomolgi* strain B vs. *P. relictum* SGS1-like** | 54.3 | 42.3 | 69.9 |
| ***P. vivax* P01 vs. *P. vivax* Sal-1** | 99.9 | 99.3 | 100 |
| ***P. vivax* P01 vs. *P. vivax* PvW1** | 99.9 | 99.4 | 100 |
| ***P. vivax* P01 vs. *P. vivax-like* Pvl01** | 96.9 | 93 | 99.1 |
| ***P. vivax* Sal-1 vs. *P. vivax* PvW1** | 99.9 | 99.4 | 100 |
| ***P. vivax* Sal-1 vs. *P. vivax-like* Pvl01** | 96.8 | 92.8 | 99.1 |
| ***P. vivax* PvW1 vs. *P. vivax-like* Pvl01** | 96.8 | 92.8 | 99.1 |
| ***P. knowlesi* strain H vs. *P. knowlesi* strain A1H1** | 100 | 100 | 100 |
| ***P. knowlesi* strain H vs. *P. knowlesi* Malayan Pk1 A** | 100 | 100 | 100 |
| ***P. knowlesi* strain A1H1 vs. *P. knowlesi* Malayan Pk1 A** | 100 | 100 | 100 |
| ***P. relictum* SGS1-like vs. *P. gallinaceum*** | 82.9 | 75.9 | 90.8 |
| ***P. falciparum* 3D7 vs. *P. falciparum* 7G8 (2019)** | 100 | 99.5 | 100 |
| ***P. reichenowi* CDC vs. *P. reichenowi* G01** | 99.4 | 98.2 | 100 |
| ***P. yoelii* 17X vs. *P. yoelii* YM** | 100 | 100 | 100 |
| ***P. ovalecurtisi* vs. *P. ovalewallikeri*** | 94.1 | 89.8 | 97.4 |

**Supplementary Table 6**: Sources for *Plasmodium* amino acid sequences used in core genome phylogenetic tree, other than *P. cynomolgi* Berok K4A7.

| ***Plasmodium*** **Species** | **PlasmoDB Link (Release 68)** | **Retrieval Date** |
|---|---|---|
| ***P. cynomolgi*** **strain M** | https://plasmodb.org/common/downloads/Current_Release/PcynomolgiM/fasta/data/PlasmoDB-68_PcynomolgiM_AnnotatedProteins.fasta | 6 March 2026 |
| ***P. cynomolgi*** **strain B (M)** | https://plasmodb.org/common/downloads/Current_Release/PcynomolgiB/fasta/data/PlasmoDB-68_PcynomolgiB_AnnotatedProteins.fasta | 7 April 2026 |
| ***P. vivax*** **Sal-1** | https://plasmodb.org/common/downloads/Current_Release/PvivaxSal1/fasta/data/PlasmoDB-68_PvivaxSal1_AnnotatedProteins.fasta | 6 March 2026 |
| ***P. vivax*** **P01** | https://plasmodb.org/common/downloads/Current_Release/PvivaxP01/fasta/data/PlasmoDB-68_PvivaxP01_AnnotatedProteins.fasta | 6 March 2026 |
| ***P. vivax*** **PvW1** | https://plasmodb.org/common/downloads/Current_Release/PvivaxPvW1/fasta/data/PlasmoDB-68_PvivaxPvW1_AnnotatedProteins.fasta | 7 April 2026 |
| ***P. vivax*****-like Pvl01** | https://plasmodb.org/common/downloads/Current_Release/Pvivax-likePvl01/fasta/data/PlasmoDB-68_Pvivax-likePvl01_AnnotatedProteins.fasta | 7 April 2026 |
| ***P. knowlesi*** **strain H** | https://plasmodb.org/common/downloads/Current_Release/PknowlesiH/fasta/data/PlasmoDB-68_PknowlesiH_AnnotatedProteins.fasta | 6 March 2026 |
| ***P. knowlesi*** **strain A1H1** | https://plasmodb.org/common/downloads/Current_Release/PknowlesiA1H1/fasta/data/PlasmoDB-68_PknowlesiA1H1_AnnotatedProteins.fasta | 7 April 2026 |
| ***P. knowlesi*** **Malayan Pk1 A** | https://plasmodb.org/common/downloads/Current_Release/PknowlesiMalayanPk1A/fasta/data/PlasmoDB-68_PknowlesiMalayanPk1A_AnnotatedProteins.fasta | 7 April 2026 |
| ***P. ovalecurtisi*** | https://plasmodb.org/common/downloads/Current_Release/PovalecurtisiGH01/fasta/data/PlasmoDB-68_PovalecurtisiGH01_AnnotatedProteins.fasta | 6 March 2026 |
| ***P. ovalewallikeri*** | https://plasmodb.org/common/downloads/Current_Release/PovalewallikeriPowCR01/fasta/data/PlasmoDB-68_PovalewallikeriPowCR01_AnnotatedProteins.fasta | 7 April 2026 |
| ***P. yoelii*** **17X** | https://plasmodb.org/common/downloads/Current_Release/Pyoeliiyoelii17X/fasta/data/PlasmoDB-68_Pyoeliiyoelii17X_AnnotatedProteins.fasta | 6 March 2026 |
| ***P. yoelii*** **YM** | https://plasmodb.org/common/downloads/Current_Release/PyoeliiyoeliiYM/fasta/data/PlasmoDB-68_PyoeliiyoeliiYM_AnnotatedProteins.fasta | 7 April 2026 |
| ***P. chabaudi*** | https://plasmodb.org/common/downloads/Current_Release/Pchabaudichabaudi/fasta/data/PlasmoDB-68_Pchabaudichabaudi_AnnotatedProteins.fasta | 6 March 2026 |
| ***P. reichenowi*** **CDC** | https://plasmodb.org/common/downloads/Current_Release/PreichenowiCDC/fasta/data/PlasmoDB-68_PreichenowiCDC_AnnotatedProteins.fasta | 6 March 2026 |
| ***P. reichenowi*** **G01** | https://plasmodb.org/common/downloads/Current_Release/PreichenowiG01/fasta/data/PlasmoDB-68_PreichenowiG01_AnnotatedProteins.fasta | 7 April 2026 |
| ***P. falciparum*** **3D7** | https://plasmodb.org/common/downloads/Current_Release/Pfalciparum3D7/fasta/data/PlasmoDB-68_Pfalciparum3D7_AnnotatedProteins.fasta | 6 March 2026 |
| ***P. falciparum*** **7G8 (2019)** | https://plasmodb.org/common/downloads/Current_Release/Pfalciparum7G8-2019/fasta/data/PlasmoDB-68_Pfalciparum7G8-2019_AnnotatedProteins.fasta | 7 April 2026 |
| ***P. gallinaceum*** | https://plasmodb.org/common/downloads/Current_Release/Pgallinaceum8A/fasta/data/PlasmoDB-68_Pgallinaceum8A_AnnotatedProteins.fasta | 6 March 2026 |
| ***P. relictum*** **SGS1-like** | https://plasmodb.org/common/downloads/Current_Release/PrelictumSGS1-like/fasta/data/PlasmoDB-68_PrelictumSGS1-like_AnnotatedProteins.fasta | 6 March 2026 |

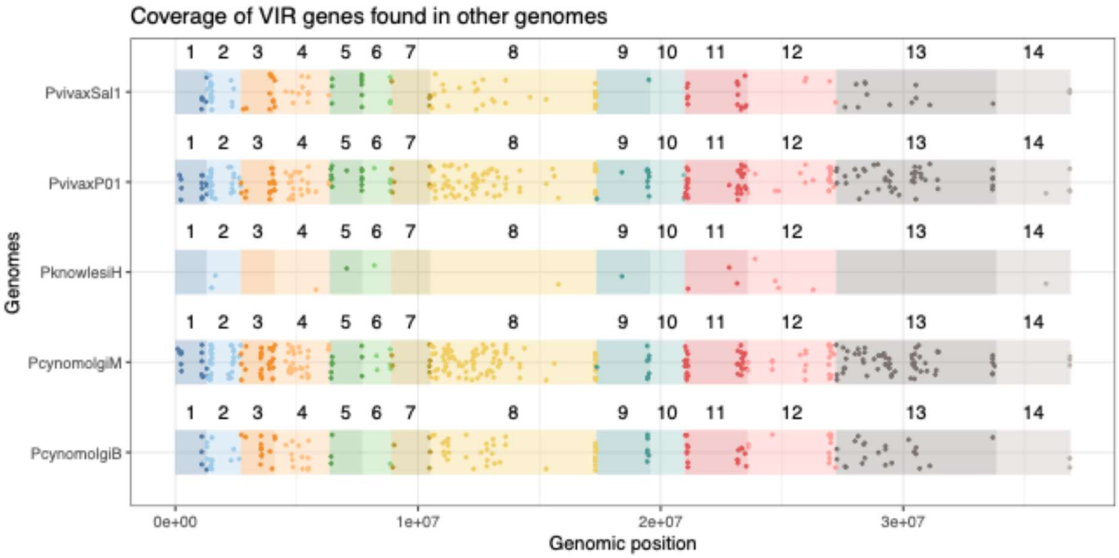


**Supplementary Figure 7.** *Vir* genes identified using blastp from various *Plasmodium* species projected on the chromosomes.
